# Learning the native-like codons with a 5’UTR and RNA secondary structure aided species-informed transformer model

**DOI:** 10.1101/2025.07.19.665668

**Authors:** Qiuyue Hu, Xiaolin Tian, Yuanning Li, Rui Zhou, Zhihao Wang, Jintao Meng, Shen Wang, Jingjing Guo, Weifeng Li, Liangzhen Zheng, Yanjie Wei

## Abstract

Efficient protein expression across heterologous hosts remains a major challenge in synthetic biology, largely due to species-specific differences in codon usage and regulatory sequence context. A key difficulty lies in reconstructing the codon landscape of the target expression system within a foreign host with a native-like codon preference. To address this, we present TransCodon, a Transformer-based deep learning model that leverages both 5’ untranslated regions (5’UTRs) and coding sequences (CDS), along with explicit species identifiers and RNA secondary structure information, to learn nuanced codon usage patterns across diverse organisms. By incorporating multisource genomic data and modeling sequence dependencies in a masked language modeling paradigm, TransCodon effectively captures both local and global determinants of codon preference. Our experiments demonstrate that integrating species-level information during training significantly improves the model’s ability to predict optimal synonymous codons when considering different evaluation metrics. More importantly it identifies native-like codons with less divergence from natural sequences compared to other methods. Besides, TransCodon could capture more low-frequency codons which are often omitted by other deep learning-based methods. The results thus indicate that TransCodon as a robust codon language model has the potential for generating native-like CDS with high translational efficiency in target hosts.

## 1 Introduction

A DNA sequence consists of four nucleotides—adenine (A), thymine (T), guanine (G), and cytosine (C). Triplets of non-overlapping nucleotides, known as codons, encode specific amino acids in a protein sequence. Multiple codons can encode the same amino acid; these are referred to as synonymous codons and are a fundamental basis for the degeneracy of the genetic code. However, the usage frequency of these synonymous codons varies significantly across species, influenced by factors such as intracellular tRNA abundance, translational dynamics, protein folding regulation, and evolutionary constraints[1, 2]. This phenomenon, referred to as codon usage bias, is a species-specific characteristic that plays a critical role in determining the efficiency of gene expression.

Codon optimization, the process of redesigning DNA sequences to align synonymous codon usage with the preferences of a target host organism, is a crucial step in heterologous protein expression. With the advent of affordable de novo DNA synthesis and advances in protein engineering[3–7], the demand for codon optimization has surged. However, the combinatorial complexity of synonymous codon arrangements makes exhaustive search approaches impractical. Traditional codon optimization methods often rely on selecting the most frequent codons or employ heuristic rules that fail to consider broader biological context[8]. Strategies focused on optimizing the Codon Adaptation Index (CAI)[9] and favoring the most common codons have shown some success in specific cases[10, 11]. However, such approaches do not guarantee effective expression and can lead to unintended consequences, including excessive cellular resource consumption, protein misfolding, ribosomal stalling, and even host toxicity[12]. Codon harmonization methods attempt to mimic natural codon usage patterns to reconstruct the codon landscape in heterologous hosts[13, 14]. Yet, their reliance on heuristic rules restricts their ability to capture deeper biological relationships, often resulting in a superficial reconstruction of codon landscapes that neglects important factors such as translational dynamics and mRNA stability.

Furthermore, many existing approaches neglect the functional role of untranslated regulatory regions, particularly the 5’ untranslated region (5’ UTR) of mRNA. Located upstream of the start codon, the 5’ UTR plays a crucial role in regulating protein expression levels[8, 15–17]. It influences mRNA stability, ribosome binding efficiency, and translation initiation. The structural features of the 5’ UTR, such as secondary structure, the sequence context around the start codon, and the ribosome binding site (RBS), are closely linked to ribosomal accessibility and overall translational efficiency. Failing to consider the 5’ UTR during codon optimization may result in inadequate modeling of translational regulation mechanisms. Additionally, the RNA secondary structure within the coding sequence (CDS) significantly affects translation[18]. Local secondary structures modulate ribosome elongation and pausing, thereby impacting protein folding and function. Strong secondary structures can cause ribosome stalling and reduce translation efficiency. Therefore, effective codon optimization should integrate both 5’ UTR and CDS structural features to better capture the dynamics of gene expression.

Recent advances in deep learning have introduced powerful alternatives for codon optimization. Neural network models have shown strong capabilities in learning complex sequence dependencies and capturing the latent “language” of codon usage[15, 19–21]. Nevertheless, current deep learning-based approaches face several limitations: they are often trained on small-scale Datasets, tailored to a single host species (e.g., *Escherichia coli*), and lack the flexibility required for broader adoption in synthetic biology. Additionally, in the context of cross-species protein expression, many models focus excessively on optimizing the CAI, which further deviates from natural codon usage patterns[20, 21]. CodonTransformer[21], a codon optimization model based on the BERT architecture, incorporates species-specific information and is trained on approximately one million sequences. However, it does not consider the regulatory effects of the 5’ UTR, and its use of the BigBird model tends to produce sequences with overly high CAI values, potentially underestimating the importance of rare codons. These limitations hinder its generalizability and biological relevance.

In this study, we present TransCodon, a novel species-aware codon optimization model based on the BERT (Bidirectional Encoder Representations from Transformers) architecture. Our model integrates multiple sequence components, including the 5’ untranslated region (5’ UTR) and coding sequence (CDS), and explicitly incorporates species-specific information and RNA secondary structure features to capture codon usage preferences across diverse organisms. By leveraging the sequence modeling capabilities of the Transformer architecture and utilizing a large-scale dataset comprising 5.5 million gene sequences from 1436 species, TransCodon effectively identifies both global and local patterns of codon usage, enabling a more accurate reconstruction of the codon landscape. This facilitates robust optimization of heterologous genes to enhance protein expression. Experimental results demonstrate that TransCodon consistently outperforms traditional methods and recent machine learning-based approaches across multiple evaluation metrics as well as identifying low-frequency codons. The TransCodon model demonstrates superior performance not only in sequence recovery rate but also across other key metrics for gene (or mRNA) sequences. These include the Codon Similarity Index (CSI), GC content, RNA secondary structure energy, and the dynamic time warping (DTW) for local codon usage fluctuations, outperforming other comparative models. Furthermore, the fitness scores calculated from codon probabilities generated by TransCodon exhibit a stronger correlation with experimentally measured protein expression levels in model organisms than those derived from other models. Notably, TransCodon effectively captures the usage of low-frequency codons, similar to patterns observed in natural gene sequences. Intriguingly, a correlation between the proportion of low-frequency codons in the predicted sequences and protein abundance an even be observed, mirroring phenomena seen in natural systems. Collectively, these findings demonstrate that TransCodon is a codon optimization model capable of generating synthetic gene sequences that closely approach their natural counterparts. Further performance improvements can be achieved through fine-tuning, validating its effectiveness and broad applicability in synthetic biology.

## 2 Results

### 2.1 A Transformer-based framework for cross-species codon optimization

Due to the inherently sequential nature of genomic data, recurrent neural networks (RNNs) and Transformer-based architectures have demonstrated remarkable efficacy in capturing the complex intrinsic features of biological sequences[22–26], thereby making them highly suitable for codon optimization tasks[20, 21, 27, 28]. In prior research, codon optimization has often been formulated as either a sequence labeling or machine translation problem. Both approaches typically use the target protein sequence as input and leverage bidirectional encoder models to generate context-aware embeddings for each amino acid, employing architectures such as bidirectional long short-term memory networks (Bi-LSTM) or bidirectional Transformer encoders[29, 30].

Given the large-scale training data used in this study, we employ a more expressive bidirectional Transformer encoder architecture, trained using a masked language modeling (MLM) approach, which has been successfully applied for pre-training protein amino acid sequences[22, 23]. A key challenge lies in defining the mapping from amino acids to codons. CodonTransformer[21] uses a composite vocabulary where each token represents an amino acid-codon pair, such as A-GCC to denote alanine encoded by the codon GCC. In contrast, our study opts for a finer-grained vocabulary based solely on nucleotides (A, T, C, and G), aligning with the design of most DNA/RNA language models[31, 32]. This choice is based on the observation that synonymous codons(different codons encoding the same amino acid) often differ by only one or two nucleotides. Therefore, instead of representing an amino acid like alanine as a generic token, such as A-UNK (an alanine residue with an unspecified codon), as in CodonTransformer, our method enables partial decoding (e.g., GC<UNK>), preserving richer sequence-level information (Figure 1a). During training, a fraction of nucleotide tokens are randomly masked with a special <mask> symbol. During inference, nucleotides that can be confidently determined are explicitly predicted, while only the uncertain ones are represented with <UNK>. This strategy enables the model to generate optimized DNA sequences conditioned on the target protein sequence with improved precision and biological fidelity. The vocabulary granularity ablation analysis is provided in Supporting Table S1 of the supplementary materials.

**Figure 1:**
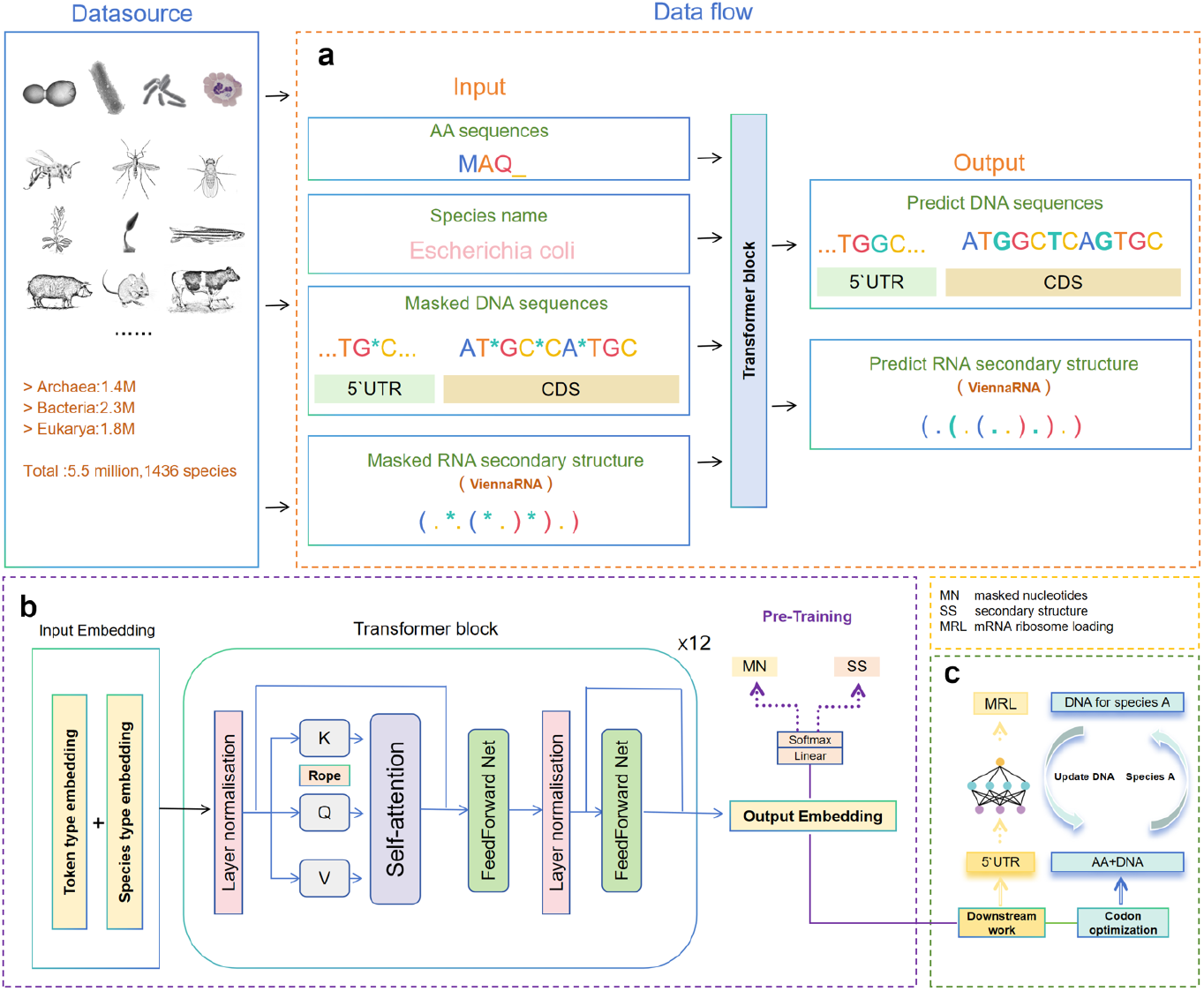
Overview of the design and applications of the codon optimization model, TransCodon. The model was trained on approximately 5.5 million samples from 1,436 species. **a. Data flow**: The model takes as input an amino acid sequence, species name, a masked DNA sequence (including both the 5’ untranslated region and the coding region), and a masked RNA secondary structure predicted by ViennaRNA. These inputs are processed through multiple Transformer layers to produce the recovered DNA sequence and RNA secondary structure. **b. Model architecture**: TransCodon adopts an encoder-only Transformer architecture. The input embeddings include token type embeddings and species identity embeddings. The training objectives comprise masked nucleotide prediction (MN) and secondary structure prediction (SS). **c. Applications**: TransCodon can be applied to downstream tasks such as 5’UTR-related MRL prediction. Additionally, it can be used for cross-species codon optimization: given a target species and an amino acid sequence, the model generates a random DNA sequence and iteratively refines it to produce an optimized DNA sequence specific to the target species.

Furthermore, to achieve species-aware optimization, we adopt a strategy inspired by the token type embedding mechanism used in Transformer architectures. Specifically, we incorporate a species embedding into the learnable token embeddings, mapping each species to a unique token type identifier. This design enables the model to explicitly capture species-specific codon usage preferences. During inference, it allows users to direct DNA optimization toward a specified target species by selecting the corresponding token type (Figure 1a). During training, RNA secondary structure information is represented in dot-bracket notation and used as a supervised training label (Figure 1a). This supervised task, similar to masked token recovery, is integrated with unsupervised masked language modeling to enable semi-supervised training. The model architecture is detailed in Figure 1a.

Our pretrained model, named TransCodon, was trained on four datasets comprising approximately 5.5 million gene sequences from 1436 species, including bacteria (41.38%), archaea (25.88%), and eukaryotes (32.75%) (Figure 1). This diverse dataset ensures robust cross-species generalization. Additionally, we constructed a holdout test dataset composed of eight representative species from across the tree of life. This test dataset was used for downstream evaluation and fine-tuning with the top 10% of high-expression genes ranked by CSI[33]. Details regarding the model architecture, training protocols, fine-tuning, and dataset preprocessing can be found in the Methods section.

### 2.2 TransCodon generates DNA sequences closely resembling native sequences

To assess the similarity between DNA sequences generated by the model and their corresponding native sequences across different species, we evaluated the generated sequences using several metrics, including codon recovery rate, CSI[33], CAI[9], GC content[34], codon frequency distribution (CFD)[35], minimum free energy (MFE)[36], and DTW[37]. Definitions of these metrics are provided in the Methods section. For a fair comparison, the test set was selected from a reserved subset of eight representative species within Dataset1 (the training set used for CodonTransformer, as described in the Methods section). These species were also included in the fine-tuning dataset for CodonTransformer. Importantly, the training and test sets were duplicated using CD-HIT[38] with a 40% sequence identity threshold, while CodonTransformer’s pretraining utilized the full dataset without duplicates.

#### 2.2.1 Codon recovery rate

Codon recovery rate is the most direct metric for assessing the similarity between generated and native sequences. As shown in Figure 2a, TransCodon outperforms uniform random choice (URC), background frequency choice (BFC), and CodonTransformer in this regard. The TransCodon-baseline model achieves an average codon recovery rate of 49.0% on the held-out test set, demonstrating its ability to preserve the inherent structure and functional features of the target sequences. Moreover, fine-tuning did not yield significant improvements over the base model, suggesting that the pretraining phase had already captured the codon usage patterns effectively. In contrast, after fine-tuning, CodonTransformer exhibited a slight improvement, reaching a codon recovery rate of 48.6%, which is comparable to that of the TransCodon-baseline model.

**Figure 2:**
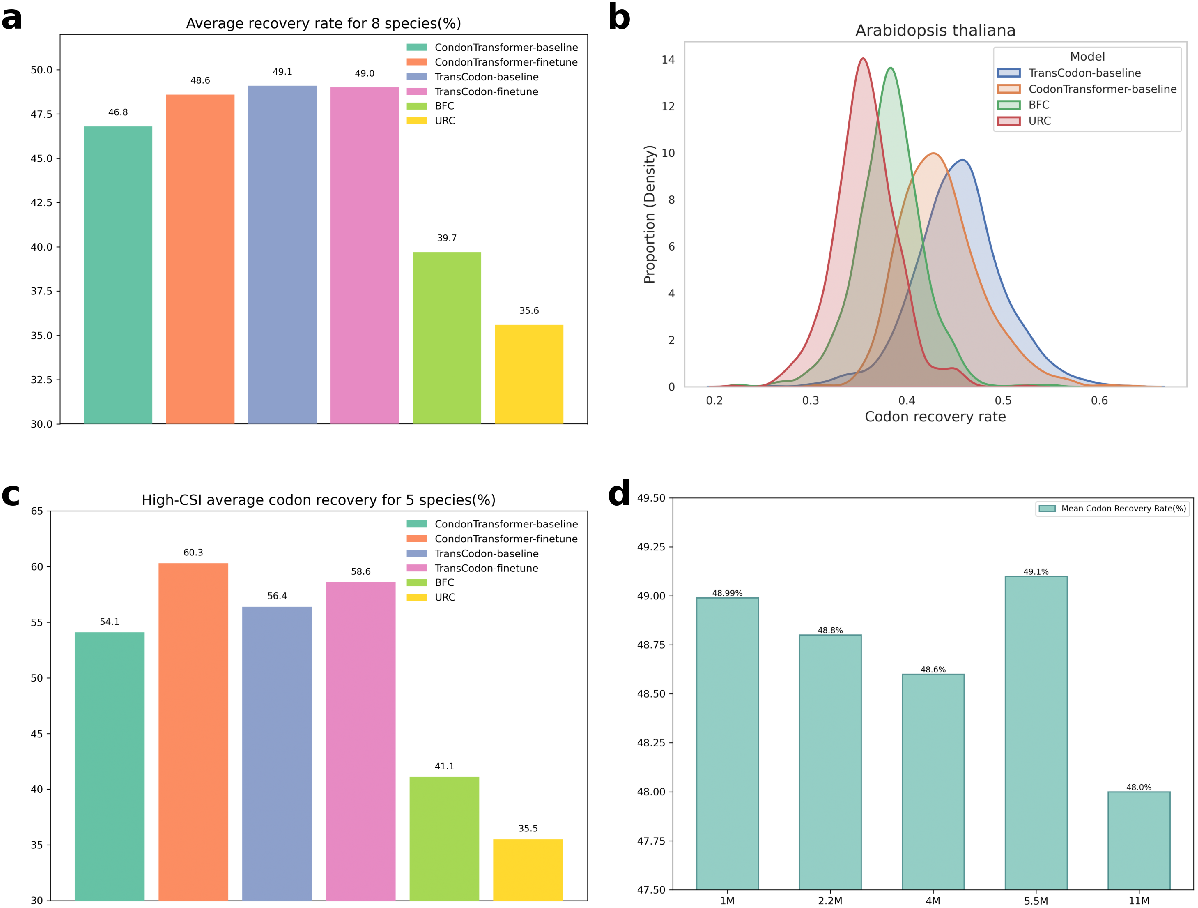
**a**. The average codon recovery rate across 8 species, with species details provided in the Methods section. **b**. Distribution of codon recovery rates for *Arabidopsis thaliana* as a representative species. **c**. Codon recovery rate for high-CSI sequences across 5 species (*Arabidopsis thaliana, Escherichia coli, Homo sapiens, Mus musculus, and Saccharomyces cerevisiae*). **d**. Impact of varying training dataset sizes on codon recovery rate, with the test set consisting of the 8 retained species from Dataset1. The training sets are as follows: 1M (Dataset1), 2.2M (Dataset1+Dataset2), 4M (Dataset1+Dataset2+Dataset3), 5.5M (Dataset1+Dataset2+Dataset4), and 11M (Dataset1+Dataset2) with an additional 8.8M bacterial data.

Following the experiments of CondonTransformer,we also analyzed the codon recovery rate of high-CSI sequences in five species. Fifty sequences per species (the top 10% in terms of CSI and included in the fine-tuning dataset) were selected for testing. As shown in Figure 2c, the TransCodon-baseline still outperforms the CodonTransformer-baseline, while CodonTransformer showed the best performance after fine-tuning, slightly surpassing TransCodon.

Although the test data originates from Dataset1, we assessed the model’s performance by progressively expanding the training set. As shown in Figure 2d, the average codon recovery rate remained stable across eight species as the training data increased from 1M to approximately 5.5M, indicating robust performance and effective assimilation of codon distribution information from diverse species. However, when the training data expanded to 11M, the codon recovery rate decreased, suggesting that 11M may exceed the effective modeling capacity of the current model parameters.

#### 2.2.2 CSI and CFD

In previous methods, when evaluating CSI/CAI and CFD metrics, it was generally assumed that higher CSI/CAI values and lower CFD values indicate better expression levels[9, 39]. While this assumption may reflect a model’s ability to capture codon usage frequencies, it also risks replicating the shortcomings of approaches that merely select the most frequent codons—potentially leading to issues such as improper protein folding. We argue that the optimal outcome is for the generated sequences to closely match the CSI/CAI and CFD distributions of natural sequences, particularly those of naturally highly expressed sequences.

Using *Arabidopsis thaliana, Mus musculus*, and *Escherichia coli general* as representative species (results for other species are provided in the supplementary materials), we compared several models based on the CSI and CFD. The Kullback-Leibler (KL)[40] divergence is a key measure of how one probability distribution diverges from a second, reference distribution—the smaller the value, the closer the distributions are. As shown in Figure 3, the BFC method, which is based purely on global codon frequencies, yields average CSI and CFD values that are closer to the natural distribution. For instance, in *Arabidopsis thaliana*, its KL divergences are 0.24 and 0.0251 for CSI and CFD, respectively. However, this method lacks deep modeling capabilities, relying purely on simple probability heuristics. In contrast, TransCodon demonstrated superior alignment with natural sequences across both metrics, achieving KL divergences of 1.29 (CSI) and 0.6174 (CFD) in *Arabidopsis thaliana*. By contrast, CodonTransformer exhibited systematic biases: its CSI values were disproportionately high, while its CFD scores deviated markedly from biological reality. The baseline model showed extreme divergence from natural data—with KL scores of 18.73 (CSI) and 2.551 (CFD) in *Arabidopsis thaliana*.Notably, fine-tuning TransCodon with high-CSI sequences elevates the CSI of predicted sequences. As expected, this also increases the KL divergences between predicted and natural distributions for both CSI and CFD metrics.

**Figure 3:**
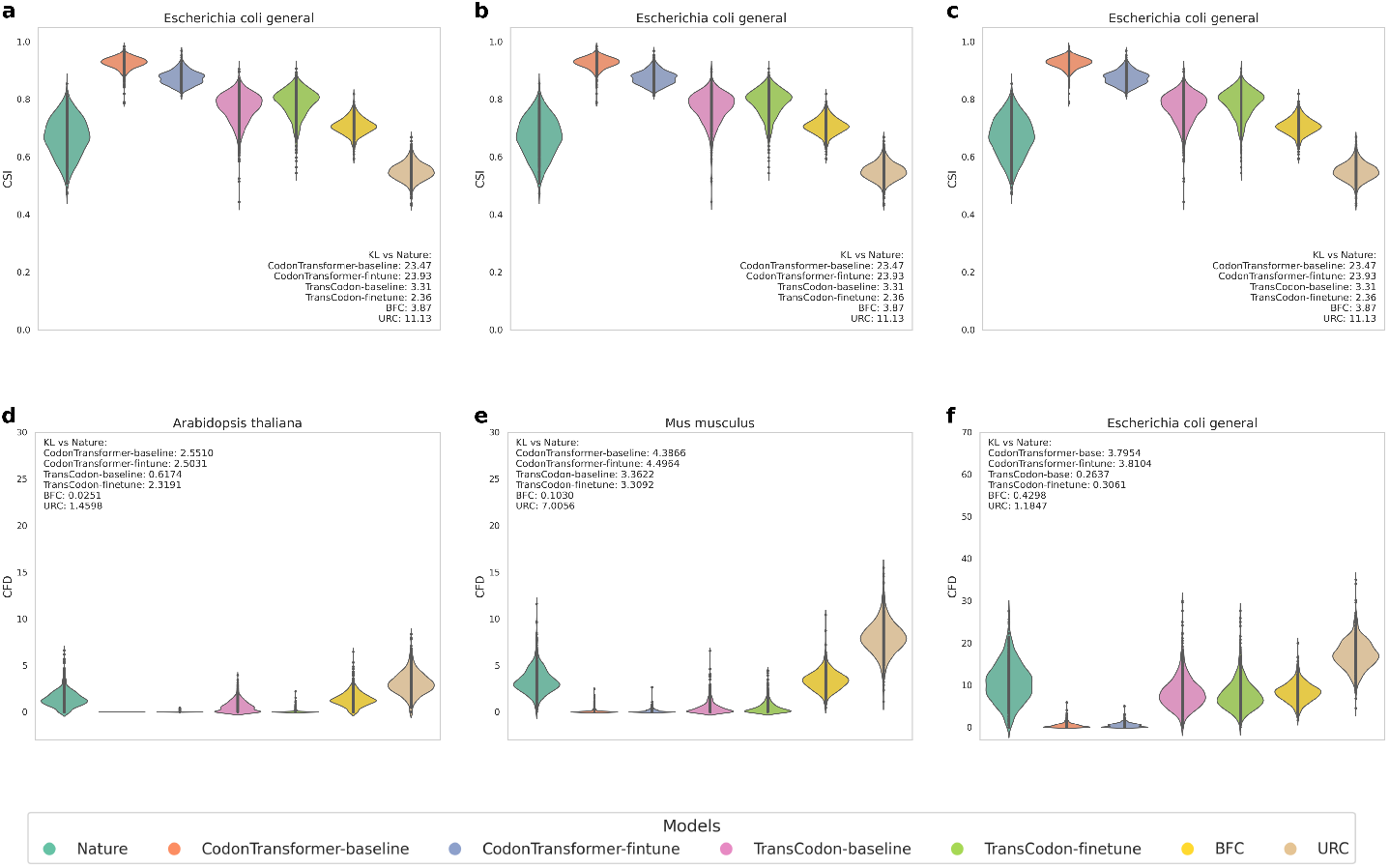
TransCodon-generated sequences closely resemble natural sequences. Using *Arabidopsis thaliana, Mus musculus*, and *Escherichia coli* as representative species (full details for all 8 species are provided in the supporting materials). **a**,**b**,**c**. In terms of CSI, both TransCodon-baseline and BFC exhibit the closest similarity to natural sequences. For *Escherichia coli*, the KL divergence between TransCodon-baseline and natural sequences is 3.31, significantly lower than CodonTransformer’s 23.47. **d**,**e**,**f**. Similarly, in CFD analysis, TransCodon-baseline and BFC most closely mimic natural distributions. Notably, fine-tuning with high-CSI genes typically increases TransCodon’s KL divergence— for example, from 0.6174 to 2.3191 in *Arabidopsis thaliana*.

#### 2.2.3 GC and MFE

GC content varies significantly across species and even among different genes within the same species[41, 42]. These variations reflect the inherent diversity of genetic sequences. Therefore, when generating sequences, it is crucial to account for this diversity rather than enforcing a fixed GC ratio. Based on this, we contend that the GC content of generated sequences should closely match the GC levels of the corresponding natural sequences, rather than adhering to a fixed range.

Accordingly, in evaluating the structural properties of generated sequences, we prioritize the consistency of minimum free energy (MFE) values with those of natural sequences, rather than merely favoring lower or higher MFE values.

Using *Drosophila melanogaster, Mus musculus*, and *Escherichia coli general* as a representative example (results for other species are provided in the supplementary materials), we compare the GC content and MFE deviation between generated and natural sequences. A smaller deviation indicates greater similarity to the natural sequence. As shown in Figure 4a,b, TransCodon demonstrates superior performance in both metrics. For instance, in *Drosophila melanogaster*, the mean differences in GC content and MFE were 2.66 and 32.42, respectively, indicating that the generated sequences closely resemble natural sequences in both GC ratio and the mRNA structural stability. In contrast, CodonTransformer exhibited the largest discrepancies in these metrics (6.71 and 65.88), suggesting a suboptimal alignment with natural patterns. Notably, TransCodon’s results were remarkably close to those of BFC (a frequency-based method), further validating that TransCodon has learned the codon frequency and natural probability distributions of the entire species, rather than merely memorizing the most frequent codons.

**Figure 4:**
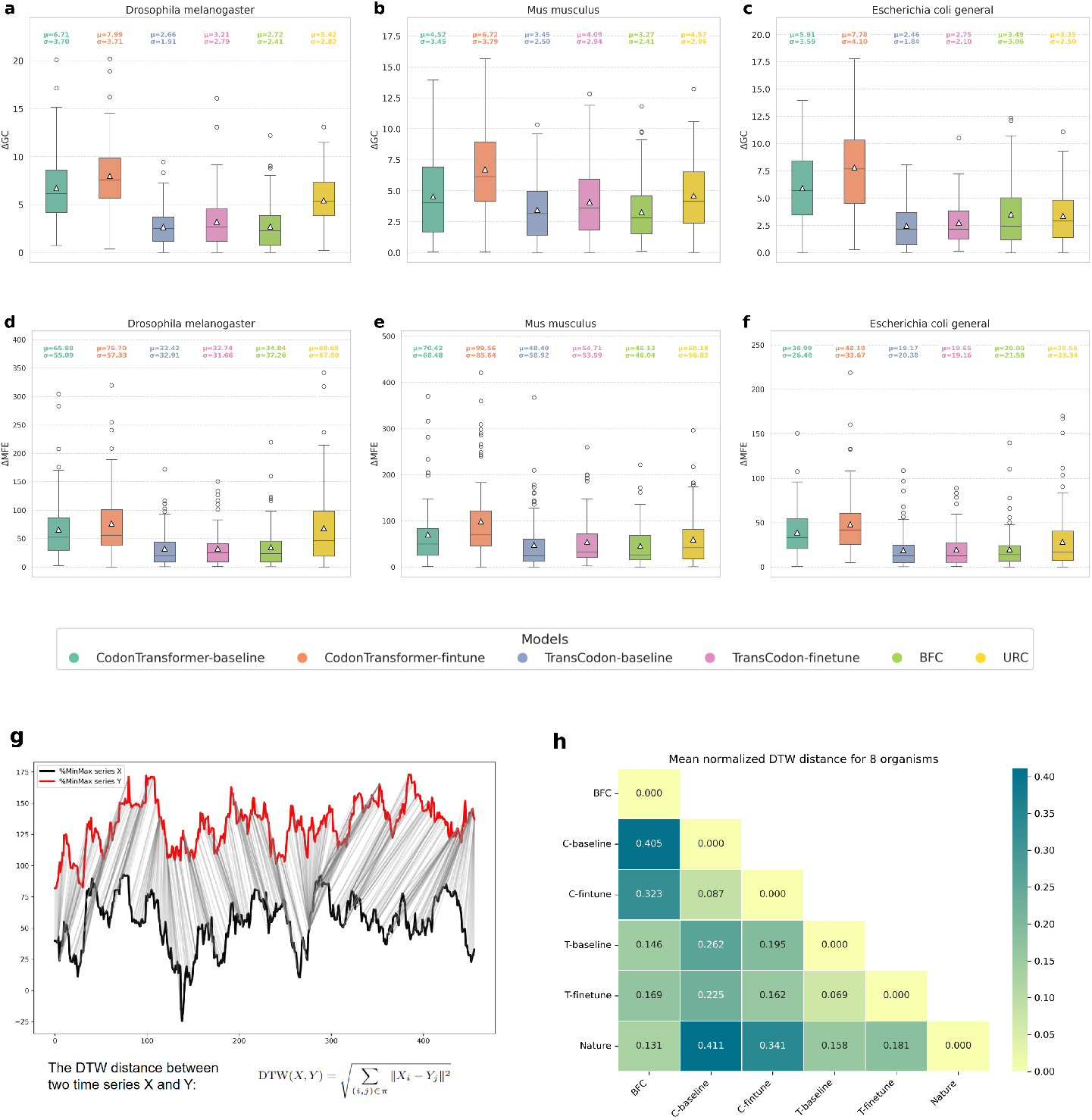
TransCodon-generated sequences closely resemble natural sequences. Using *Drosophila melanogaster, Mus musculus*, and *Escherichia coli* as representative species: **a**,**b**,**c**. In terms of GC content, both TransCodon-baseline and BFC exhibited the closest alignment with natural sequences. For example, in *Escherichia coli*, the average GC content difference between TransCodon-baseline and natural sequences was 2.46, significantly lower than the 5.91 observed in CodonTransformer. **d**,**e**,**f**. Similarly, regarding Minimum Free Energy (MFE), both TransCodon-baseline and BFC closely approximated the MFE values of natural sequences. Notably, after fine-tuning with high-CSI genes, the MFE difference for CodonTransformer increased significantly, whereas TransCodon showed minimal changes. **g**. Schematic diagram of %MinMax for the gene *yahG (Escherichia coli)* between natural and TransCodon-generated sequences. The DTW algorithm calculates the minimum distance between two %MinMax sequences by finding matching positions (see “Methods” for details).**h**. The average DTW values across 8 retained test species. C-baseline and T-baseline denote CodonTransformer-baseline and TransCodon-baseline, respectively. Results show that sequences generated by the BFC method and TransCodon have the smallest DTW distances to natural sequences, with values of 0.131 and 0.158, respectively.

#### 2.2.4 %MinMax and DTW

The %MinMax[43] metric captures local fluctuations in codon usage, offering detailed insights into patterns that influence translation efficiency and mRNA stability. However, existing methods often neglect these localized dynamics. Figure 4c illustrates two %MinMax profiles, with their similarity assessed using the Dynamic Time Warping (DTW) algorithm, which calculates the minimum alignment distance. Ideally, the %MinMax profile of generated sequences should closely match that of the corresponding natural sequences, preserving both global codon usage trends and local sequence features.

As shown in Figure 4d, comparative analysis of different models reveals that the %MinMax curves of sequences generated by TransCodon exhibit smaller DTW distances compared to natural sequences. The average DTW distance of TransCodon across 8 species is 0.158, significantly lower than the 0.341 and 0.411 values observed for CodonTransformer. Notably, in the original CodonTransformer paper, the DTW calculation was divided by 2. Additionally, its high DTW value is closely tied to the very high mean of the generated MinMax sequences, which is also associated with the excessively high CAI of its sequences. TransCodon, on the other hand, shows a smaller DTW distance of 0.146 when compared to the BFC method. As expected, fine-tuning TransCodon results in a slight increase in DTW distance, likely due to the use of high-CSI genes for fine-tuning. Although the BFC method demonstrates the smallest DTW distance to natural sequences, it relies solely on probabilistic random selection. In contrast, TransCodon-generated sequences align more closely with natural distributions and exhibit higher codon recovery rates.

In summary, the results across multiple evaluation metrics demonstrate that TransCodon effectively captures the natural codon distribution of different species and generates sequences that closely resemble natural DNA.

### 2.3 TransCodon exhibits robust effectiveness for protein abundance correlation in zero-shot scenarios

Beyond its ability to design sequences for heterologous expression systems, TransCodon also shows a certain level of correlation with experimentally determined protein abundance[44]. Previous studies have demonstrated that mRNA levels are strongly correlated with protein abundance, whereas the CAI shows a weaker correlation with protein abundance[39, 45]. In this context, the fitness score of a sequence evaluated by TransCodon reflects the likelihood of observing the corresponding DNA (or mRNA) sequence given the protein sequence. Therefore, it can be used as an index for predicting protein abundance.

We retrieved protein expression data for various genes from species including *Escherichia coli, Saccharomyces cerevisiae, Arabidopsis thaliana,Mus musculus,Drosophila melanogaster*, and *Homo sapiens* using the PaxDb database[44]. By using the native DNA sequences as input, we evaluated the correlation between protein abundance and the fitness score computed by different models. Details of the fitness score calculation can be found in the Methods section. As shown in Figure 5a,b, TransCodon consistently achieves the highest Spearman and Pearson correlation coefficients across all test species in the zero-shot setting, indicating that the model has effectively captured deep and biologically meaningful patterns.

**Figure 5:**
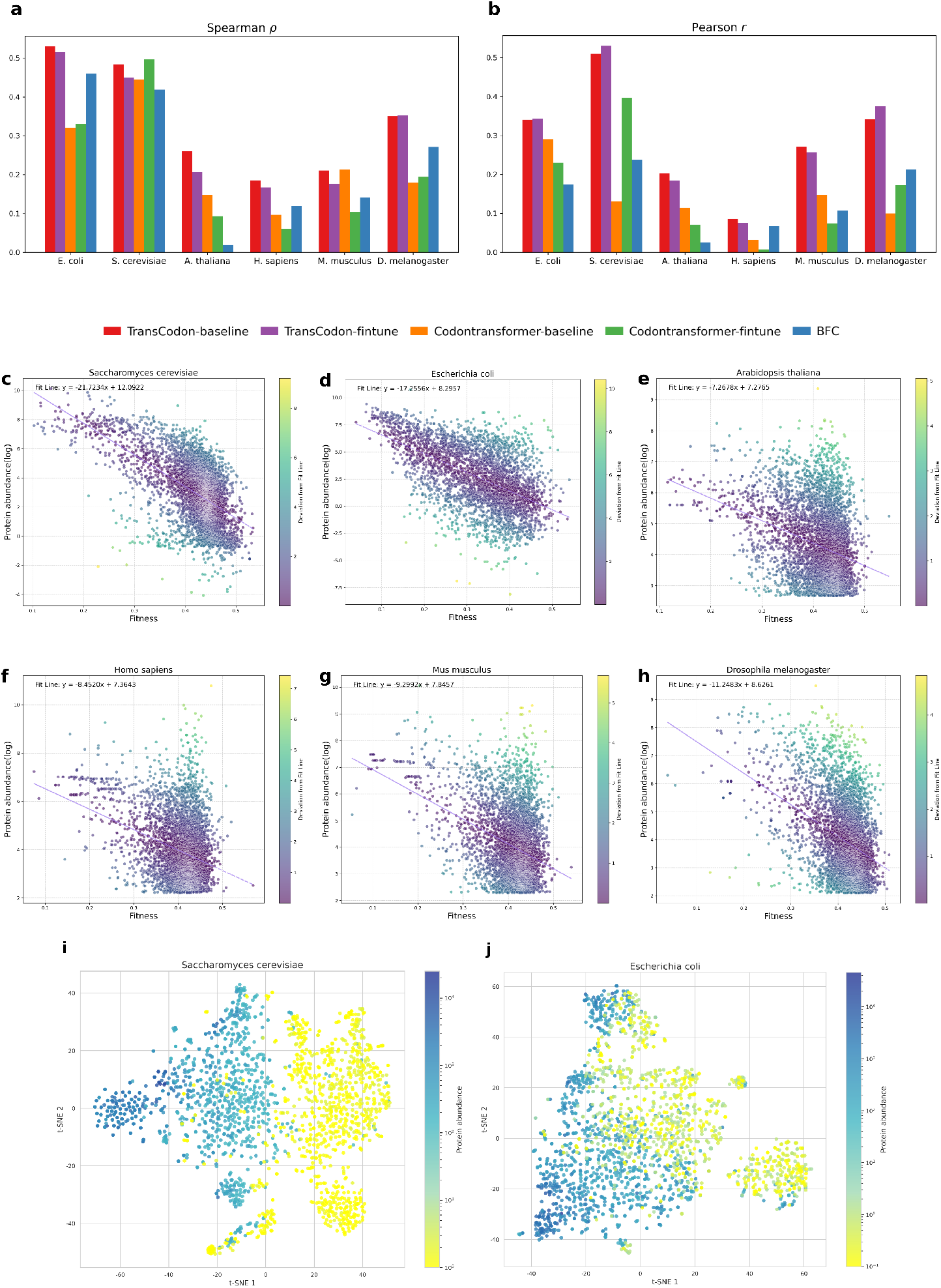
The correlation between model-predicted fitness score and protein abundance of different genes from *Escherichia coli, Saccharomyces cerevisiae, Arabidopsis thaliana, Homo sapiens, Mus musculus*, and *Drosophila melanogaster*. **a**,**b**. The correlation between protein abundance and model fitness across six species using various methods, including TransCodon, CodonTransformer, URC, and BFC. **c-h**. The relationship between protein abundance and model fitness across six species. The y-axis represents the log-transformed protein abundance, and the x-axis denotes the model’s fitness score. The color bar indicates the deviation of different points from the fit line: the greater the deviation, the lighter the color. Notably, *Escherichia coli* and *Saccharomyces cerevisiae* exhibit a distinct trend, where lower fitness scores are associated with higher protein abundance. **i**,**j**. Dimensionality reduction of embeddings for the top 1,000 and bottom 1,000 protein abundance sequences for *Escherichia coli* and *Saccharomyces cerevisiae* using TransCodon.

The high correlation observed in prokaryotes such as *Escherichia coli* can be attributed to the relatively simple regulatory mechanisms[46], where cis- and trans-acting elements exert certain influence on gene expression. As a result, codon usage remains a more dominant factor, making expression levels more predictable based solely on sequence features.

In contrast, gene expression in complex eukaryotic organisms is influenced by a wide range of regulatory mechanisms[47], including enhancers, silencers, chromatin accessibility, and post-transcriptional regulation. These layers of regulation introduce considerable biological variability, which can diminish the direct correlation between codon-level features and protein abundance — particularly in Pearson correlation, which is sensitive to linear relationships. Notably, our model continues to perform well on *Saccharomyces cerevisiae*, a relatively simple unicellular eukaryote. This result further underscores TransCodon’s robustness in capturing codon-related fitness across diverse biological contexts.

Furthermore, we examined the relationship between protein abundance and model fitness scores across different sequences within each species. The y-axis represents the log-transformed protein abundance, as determined by mass spectrometry, while the x-axis corresponds to the model’s fitness score. As illustrated in Figure 5c, all six species exhibit a consistent trend, where lower fitness scores correlate with higher protein abundance. This pattern is especially pronounced in *Escherichia coli* and *Saccharomyces cerevisiae*, and aligns with the computed Spearman correlation coefficients.

Specifically, we applied t-SNE to reduce the dimensionality of the embeddings for the top 1,000 and bottom 1,000 protein abundance sequences. As shown in Figure 5d,e, the embeddings corresponding to high and low protein abundance show clear separation for both *Saccharomyces cerevisiae* and *Escherichia coli* (see Supplementary Materials for results from other species). This observation further supports the model’s ability to capture meaningful biological information.

### 2.4 5’ UTR-Based downstream prediction of mean ribosome load (MRL)

Ribosome load refers to the number of ribosomes actively engaged in translating a given mRNA molecule at a specific time [48]. The Mean Ribosome Load (MRL) is the average number of ribosomes translating a particular mRNA and serves as a critical indicator of translational efficiency. MRL is influenced by various factors, including the sequence of the 5’ untranslated region (5’ UTR) and the secondary structure of the mRNA. UTR-LM[15], a BERT-based pre-trained model specifically trained on 5’ UTR sequences, has demonstrated superior performance in predicting mean ribosome load (MRL) compared to other methods, such as Optimus[49], FramePool[50], MTTrans[51], RNABERT[52], and RNA-FM[31]. Our model, TransCodon, also integrates 5’ UTR data during training on Dataset2 and Dataset3, prompting us to evaluate its effectiveness for 5’ UTR-related downstream tasks.

We utilized the U1, U2, *ψ*1, and *ψ*2 libraries provided by UTR-LM—synthetic sub-libraries containing 50-nucleotide 5^*′*^ UTR sequences—for Mean Ribosome Load (MRL) prediction. The data were split using a rank-based strategy[15], with each library comprising approximately 260,000 sequences for training and around 20,000 sequences for testing. As shown in Figure 1c, the final output embedding is obtained after passing through multiple Transformer layers. An MRL prediction head, implemented as a simple multilayer perceptron (MLP) with one hidden layer, is added on top of the TransCodon-baseline model. Following the UTR-LM approach[15], we perform full-parameter fine-tuning for MRL prediction.

Experimental results shown in Figure 6 demonstrate that, using the same simple MLP prediction head, TransCodon outperforms UTR-LM[15] across Spearman and Pearson correlation metrics, as well as RMSE and MAE. Furthermore, it significantly surpasses RNA-BERT[52] and RNA-FM[31] in performance. These results indicate that TransCodon excels not only in codon optimization but also in downstream tasks related to the 5’ UTR.

**Figure 6:**
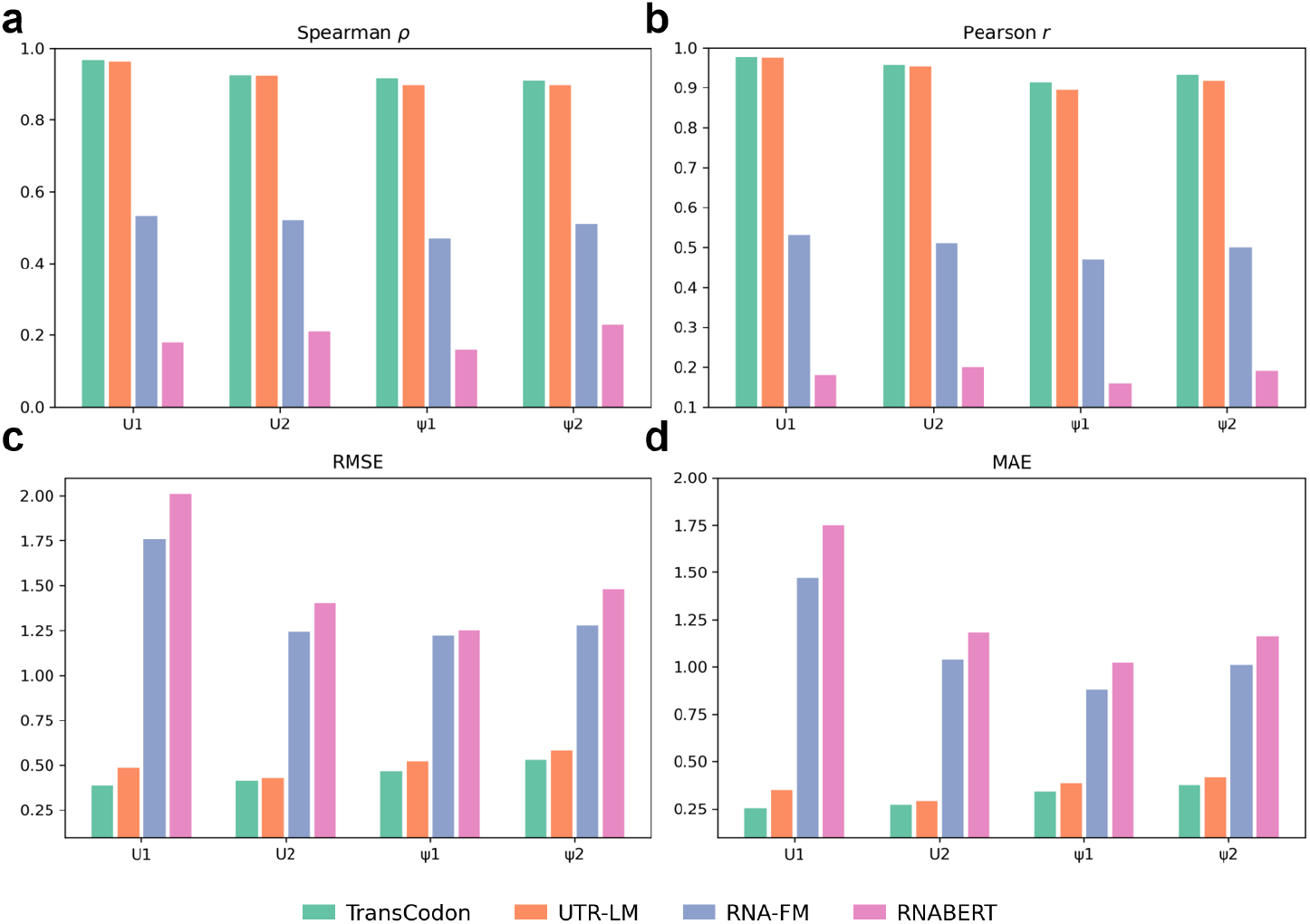
**a, b, c, d**. Performance comparison of MRL prediction across different methods, evaluated using Spearman and Pearson correlation coefficients, RMSE, and MAE. The results for *RNABERT, RNA-FM*, and *UTR-LM* are taken directly from the original UTR-LM publication.

## 3 Discusstion

### 3.1 Key features contribute to more biological meaningful fitness scores

To gain deeper insights into the factors influencing model performance,we conducted a series of ablation studies to evaluate the effects of training data volume, RNA secondary structure, and the role of the 5’UTR region. The results also warrant further investigation.

In the development of large models, both training data size and model parameters are critical for enhancing performance. Since TransCodon incorporates species labels, the DNA data for each species is limited, and expanding the training set introduces new codon distributions from additional species. This raises the question of how these new species affect the codon landscape of existing species. With a fixed test set, increasing the training data volume results in minimal changes to performance metrics, yet the correlation between model fitness and protein abundance is significantly influenced by the type of training data. From Figure S1, we observe that for eukaryotes such as *Drosophila melanogaster, Homo sapiens*, and *Mus musculus*, the correlation decreased as training data increased from 1M to 2.2M. This is likely due to the inclusion of archaea (Dataset2), whose codon usage deviates from that of eukaryotes. However, when the data size reached 5.5M (including fungal data), the correlation peaked. For microorganisms like *Escherichia coli* and *Saccharomyces cerevisiae*, adding archaea and bacterial datasets improved correlation, but a slight decrease was observed with the inclusion of fungal data. This suggests that model performance is influenced by both the scale and the composition of the training data. Expanding the training set from 1M to 5.5M enhances the model’s ability to capture codon usage preferences across species, with species diversity playing a crucial role in model generalization. Whether increasing model parameters to accommodate larger, more complex datasets will further enhance performance remains an area for future exploration.

RNA secondary structure plays a crucial role in transcription, particularly in the regions near the ribosome binding site (RBS) and the first few codons. Exposed RNA secondary structures near the RBS can enhance transcription efficiency by facilitating the binding of transcription factors or RNA polymerase. Furthermore, the secondary structure of the first few codons has been shown to influence the initiation and regulation of transcription[53, 54]. However, a unified and quantitative description of how RNA secondary structure affects transcription remains elusive. While this study does not explicitly explore the detailed patterns between RNA secondary structure and transcription, we acknowledge that its inclusion can undoubtedly improve model performance, particularly in terms of capturing more native-like properties. Although meaningful patterns may be challenging to identify in this context, the presence of RNA secondary structure in the model may provide valuable insights and help improve predictions related to transcriptional behavior. While the inclusion of RNA secondary structure information did not significantly impact codon recovery, with an average codon recovery rate of around 49%, it revealed a notable pattern in the correlation between fitness and protein abundance. The addition of RNA secondary structure improved the Spearman correlation across all five species, particularly for *Arabidopsis thaliana*, where the correlation increased from 0.19 to 0.34 (see Figure S3), suggesting its potential for providing deeper biological insights. Detailed results of all ablation experiments are provided in the supplementary materials.

Additionally, incorporating 5’ UTR information during pretraining has proven effective in enhancing model performance. The 5’ untranslated region (UTR) plays a key role in regulating gene expression by influencing the binding of regulatory proteins and ribosomes, thereby affecting translation initiation [55, 56]. Including 5’ UTR data during pretraining allows the model to better capture these regulatory elements, optimizing codon selection in a way that reflects native biological processes. As shown in Figure S2, the codon recovery rate increased from 58% to 59%, and the model’s fitness correlation with protein abundance also improved to some extent. However, omitting 5’ UTR information during inference can hamper the model’s performance, leading to a decrease in both codon recovery rate and correlation, as the model may require the same input format during inference as it did during training to maintain optimal functionality. The inclusion of 5’ UTR information during pretraining not only enhances model performance but also helps preserve the native-like properties of codons. Moreover, it highlights the importance of maintaining consistent input data across both the training and inference phases.

### 3.2 TransCodon generates native-like low-frequency codons

Many existing codon optimization tools primarily focus on maximizing the CAI[20, 21], which may inadvertently lead to issues such as protein misfolding. For multi-domain proteins, some studies[14, 57, 58] suggest that regions between the N-terminal and downstream domains are often enriched in low-frequency codons, potentially slowing translation to facilitate proper folding. However, this is not a universal principle.

For the six species mentioned above, using sequences obtained from PaxDb[44], we first sorted the DNA sequences according to protein abundance. Then, we calculated the percentage of low-frequency codons (defined as codons with a frequency of less than 0.3, as detailed in the Methods section) for each sequence. Our goal was to examine whether natural sequences exhibit any pattern in the distribution of low-frequency codons as protein abundance changes—specifically, whether highly expressed proteins tend to have a lower proportion of low-frequency codons[45]. We also sought to investigate whether TransCodon, compared to other methods, observes the same trend.

Figure 7 (a-f) shows the enrichment plots of the proportion of low-frequency codons versus protein abundance for *Escherichia coli* using different methods and natural sequences. It is evident that, in natural sequences, as protein abundance decreases, the proportion of low-frequency codons tends to increase. Figure 7 g presents the average proportion of low-frequency codons in different protein abundance ranges for the various methods and natural sequences. The line for natural sequences shows a clear upward trend. In DNA prediction methods, BFC randomly selects codons based on probabilities; although the average value is close to that of the natural sequence, the line shows almost no fluctuation. Similarly, CodonTransformer, due to its high CSI values for predicted sequences, has a lower average line, which is also relatively flat, indicating that it has not learned the pattern of low-frequency codon usage across sequences with varying protein abundances. In contrast, for TransCodon, while the average value is slightly lower than that of the natural sequence, we observe a clear upward trend in the line, which matches well with the natural sequence, suggesting that TransCodon has learned the low-frequency codon usage pattern associated with different protein abundance levels, similar to the natural sequence. The same conclusion holds for *Saccharomyces cerevisiae*, as shown in Figures 7 (h-n). Information for all six species can be found in the Supplementary Material.

**Figure 7:**
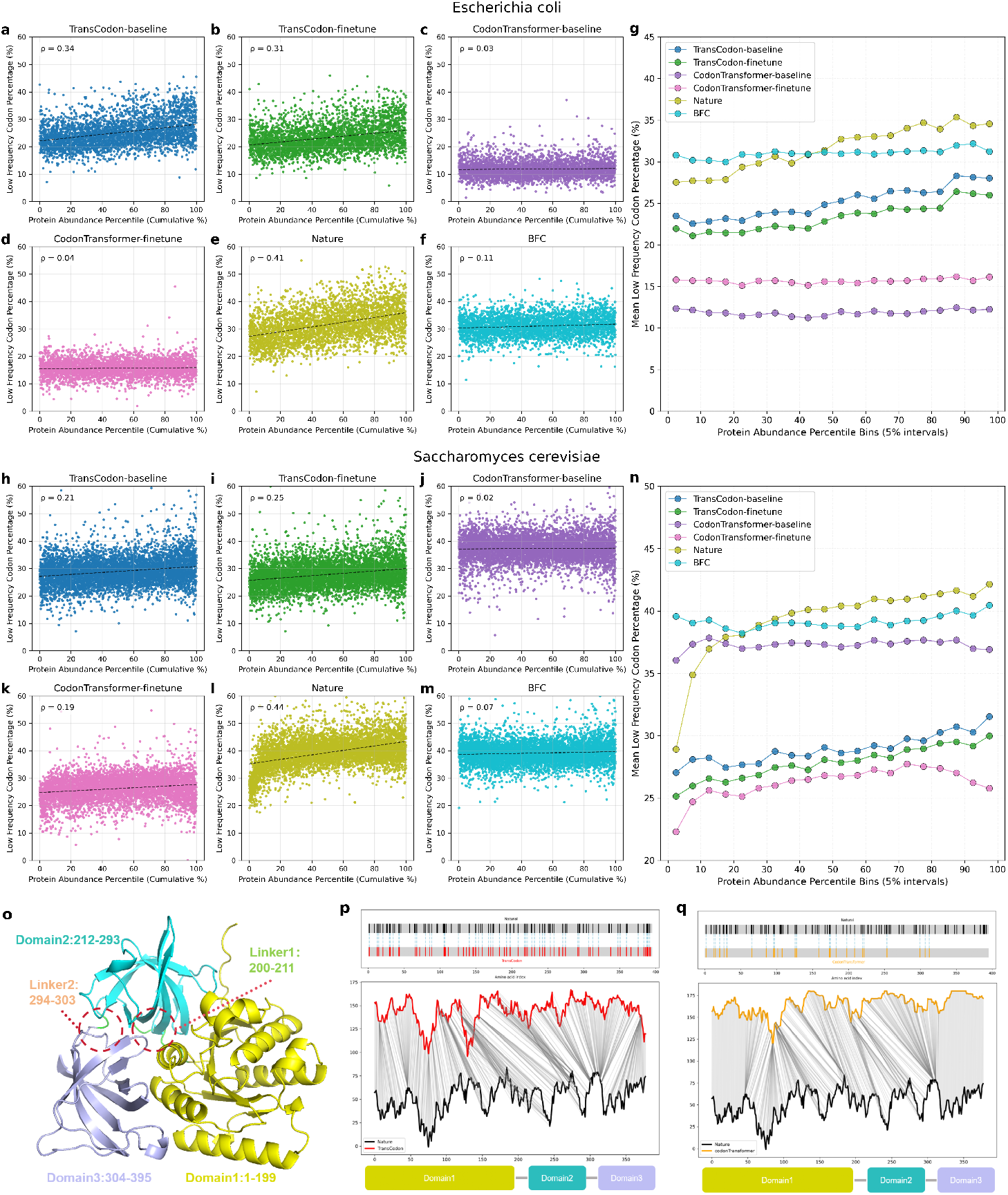
TransCodon captures the distribution of low-frequency codons. **a-g**. For *Escherichia coli*, panels **(a-f)** display enrichment plots comparing the proportion of low-frequency codons to protein abundance across different methods versus the native sequence. The x-axis represents the ranked protein abundance of sequences (with 100% indicating the lowest protein abundance), and the y-axis shows the proportion of low-frequency codons in each sequence. In panel **(g)**, the x-axis represents protein abundance percentile bins (in 5% intervals). For example, [0,5) corresponds to sequences with protein abundance in the top 5%, [95,100) represents sequences with protein abundance in the bottom 5%. The y-axis depicts the average proportion of low-frequency codons within each bin. **h-n**. Illustrate the relationship between protein abundance and low-frequency codon usage in *Saccharomyces cerevisiae*. **o**. The 3D structure of the *tufA* protein, which consists of three structural domains and two linker regions. The numbers indicate the amino acid sequence positions. **p**,**q**. In these panels, black represents the natural sequence, red represents the TransCodon-generated sequence, and orange represents the CodonTransformer-generated sequence. The top panel displays the distribution of low-frequency codons, with blue dashed lines marking the matches. The middle panel shows the %MinMax profiles, and the bottom panel provides a summary of the structural domains.

Additionally, we examined the first highly expressed multi-domain protein (*tufA*, Elongation Factor Tu) in *Escherichia coli* [44]. *tufA* is involved in the GTP-dependent binding of aminoacyl-tRNA to the A-site of ribosomes and plays a crucial role in bacterial growth as well as in the cell’s response to nutrient deprivation [59]. As shown in Figure 7o, this gene encodes a protein with three domains and two inter-domain linkers. Defining low-frequency codons as those with a usage frequency below 0.3, we identified 85 low-frequency codons out of 395 total codons in the *tufA* coding sequence, yielding a proportion of 0.2152. In the two inter-domain regions, 5 out of 22 codons were low-frequency, corresponding to a proportion of 0.227, which is comparable to the overall frequency.

What we aim to investigate is whether TransCodon, given only the amino acid sequence as input, can effectively learn which regions should incorporate low-frequency codons to slow down translation and promote stable protein folding. The TransCodon-generated DNA sequence achieves a codon-level recovery rate of 72.91% compared to the natural CDS *tufA* sequence. As shown in Figure 7p, the top panel illustrates the distribution of low-frequency codons in both the natural and TransCodon-generated sequences, the middle panel presents the %MinMax profiles of the *tufA* gene, and the bottom panel summarizes the composition of each structural domain. Notably, TransCodon achieves a 63% match rate with the natural sequence in terms of low-frequency codon positions, significantly outperforming CodonTransformer, which only achieves 28%. When examining the %MinMax sequence plot in the mid-range (indices 0–100), TransCodon demonstrates strong alignment with the natural sequence’s %MinMax pattern. Between indices 100–300, although direct matches are absent, TransCodon shows a slight lag compared to the natural sequence, yet retains a close morphological resemblance. In contrast, CodonTransformer’s alignment becomes progressively harder to discern after index 100, which is consistent with its lower matching rate for low-frequency codons. This suggests that TransCodon effectively identifies regions where low-frequency codons should be incorporated, implying that it has implicitly learned structural and folding-related biological insights.

The results indicate a correlation between the proportion of low-frequency codons and protein abundance[45, 60]. Highly abundant proteins, translated more frequently, typically exhibit a lower proportion of low-frequency codons, thereby optimizing translation efficiency. In contrast, less abundant proteins, expressed at lower levels, can afford to incorporate more rare codons without significantly impacting translation efficiency. This correlation is primarily driven by evolutionary pressures for optimal ribosome utilization in fast-growing cells, where ribosomes are a limited resource. Highly expressed proteins utilize faster codons to minimize ribosomal load and enhance translation rates, while proteins with lower expression levels are less constrained by translation speed. Additionally, factors such as tRNA availability and the need for specialized protein folding mechanisms can influence the use of low-frequency codons. While protein abundance generally correlates with a reduction in low-frequency codons, exceptions may occur due to variations in cellular context and evolutionary factors. Despite this biological nuance, methods like BFC and CodonTransformer struggle to effectively capture this relationship (Figure 7). Models that prioritize high codon CSI, particularly in sequence modeling tasks such as masked recovery, tend to minimize loss by favoring high-frequency codons[20, 21]. This results in overlooking the role of low-frequency or rare codons, a phenomenon especially evident in CodonTransformer.

In contrast, TransCodon, with its full attention mechanism, excels in capturing the global patterns of codon usage. Additionally, by incorporating RNA secondary structure information, low-frequency or rare codons may contribute crucial secondary structure elements, thus enhancing their functional significance. While the training data for the six species tested does not include the 5’ UTR region, ablation experiments have already demonstrated its importance. Therefore, considering the inclusion of the 5’ UTR region in future widespread biological modeling efforts is a promising direction to explore.

## 4 Conclusion

TransCodon establishes a transformative paradigm for codon optimization through its species-informed transformer architecture that integrates 5’UTR regulatory contexts, coding sequences, and RNA secondary structural features. By leveraging multi-organism genomic patterns across many species, the model achieves unprecedented biological fidelity—generating sequences with 49.1% codon recovery while maintaining native-like distributions of codon adaptation (KL divergence =1.29 for CSI) and low-frequency codon positioning (63% retention in structural domain interfaces). Crucially, TransCodon resolves the historical trade-off between expression optimization and sequence divergence: it simultaneously enhances protein abundance correlation (Spearman *ρ* = 0.53 in *Escherichia coli*) and preserves translational dynamics critical for proper folding. The framework’s versatility is further demonstrated through state-of-the-art performance in 5’UTR-mediated ribosome load prediction, showcasing its capacity to model end-to-end translational regulation. With openly available model weights and optimization pipelines, TransCodon provides a foundational tool for synthetic biology that bridges species-specific codon landscapes while enabling future extensions into chromatin-aware eukaryotic optimization and de novo protein design workflows.

## 5 Methods

### 5.1 Datasets

The pretraining data are compiled from four distinct datasets. Dataset 1, sourced from CodonTransformer, consists of approximately 1 million protein-coding sequences from 164 species, with bacterial, archaeal, and eukaryotic sequences representing 56.1%, 2.5%, and 41.4%, respectively. Datasets 2–4 include sequences retrieved from the NCBI database[61], encompassing approximately 1.4 million archaeal sequences (Dataset 2), 1.8 million bacterial sequences (Dataset 3), and 1.2 million eukaryotic sequences (Dataset 4). Importantly, Datasets 2 and 3 also contain 5’ untranslated regions (5’ UTRs), defined as the 100 nucleotides upstream of the start codon.

To rigorously assess both the model’s expressiveness and generalization ability, and to ensure a fair comparison with CodonTransformer, we constructed a holdout test set comprising 8 representative species from Dataset 1: *Escherichia coli O157:H7 str. Sakai, Escherichia coli str. K-12 substr. MG1655, Homo sapiens, Arabidopsis thaliana, Saccharomyces cerevisiae, Escherichia coli (general), Drosophila melanogaster*, and *Mus musculus*. All eight species are among the 15 species used for fine-tuning in CodonTransformer. To construct the test set, we applied CD-HIT[38] clustering at a 40% amino acid sequence similarity threshold and randomly selected 15% of the clusters from each species (up to a maximum of 1,000 sequences). Following filtering and preprocessing, the final held-out dataset consists of approximately 6,400 sequences.

The fine-tuning dataset was derived from the training data of the aforementioned 8 species, specifically selecting the top 10% of genes with the highest CSI scores within each species. All preprocessing steps ensured the removal of sequence similarity between the training and test datasets. Consequently, the final fine-tuning dataset comprises approximately 18,000 sequences.

**Table 1:** Dataset Summary.

| Dataset | Species | Volume | With 5'UTR | Source |
| --- | --- | --- | --- | --- |
| Dataset1<sup>1</sup> | 164 | 1,001,197 | No | CodonTransformer |
| Dataset2 | 538 | 1,407,346 | 100bp | Archaea |
| Dataset3 | 584 | 1,728,836 | 100bp | Bacteria |
| Dataset4 | 154 | 1,398,475 | No | Eukarya |

<sup>1</sup> Dataset1 contains Bacteria (561,625), Eukarya (414,445), and Archaea (25,127).

### 5.2 Model details

#### 5.2.1 Model architecture

The model architecture is inspired by the ESM family of language models and consists of three primary components: a learnable embedding layer, a stack of Transformer encoder layers, and a language modeling head. The vocabulary comprises 10 tokens, including the four nucleotides (A, T, C, G), along with special tokens to represent sequence start (<cls>), end (<eos>), padding (<pad>), unknown nucleotides (<unk>), masking (<mask>), and a separator (<sep>) to distinguish between the 5’UTR and coding regions. Input token sequences of length T are first embedded into a 768-dimensional latent space, resulting in a [*T*, 768] representation. This embedding is then processed through a Transformer encoder stack. Each self-attention layer employs multi-head scaled dot-product attention, where queries (Q), keys (K), and values (V) are projected from the inputs via learnable linear transformations. Our model uses 12 attention heads with a hidden size of 64 per head (*d*_*k*_ = 64), ensuring that each head captures distinct sequence features. To facilitate sequential modeling, rotary positional embeddings are applied instead of absolute position embeddings. Each Transformer block also includes residual connections, a feedforward neural network with a hidden dimension of 3072, and layer normalization. Furthermore, we employ pre-layer normalization and disable dropout to enhance training stability. After passing through 12 Transformer layers, the model outputs token representations of shape [*T*, 768]. These representations are then fed into the language modeling head, which consists of a feedforward network, a layer normalization step, and a matrix multiplication with the transposed embedding matrix. The head projects the 768-dimensional features to the vocabulary size, producing logits that, after applying the softmax function, yield uncalibrated probability distributions over the tokens. The model is trained using a masked language modeling (MLM) objective. For each sequence *x*, a subset of positions *M* is randomly sampled for masking, and the model is trained to minimize the negative log-likelihood of the true tokens at masked positions, given the masked input sequence.

#### 5.2.2 Model input

The model takes both DNA sequences and their corresponding species IDs as input. To enable species-aware optimization, we incorporate a token type embedding mechanism inspired by the Transformer architecture, where each token embedding is combined with a learnable species embedding.

### 5.3 Training Objectives and Model Training

The training process of our model consists of two main stages: **pretraining** and **fine-tuning**, both utilizing a dynamic masking strategy. In each training batch, 25% of input tokens are randomly selected for masking: 80% of these are replaced with a special <mask> token, 10% are substituted with a random nucleotide, and the remaining 10% remain unchanged. The model supports sequences of up to 2048 tokens. For sequences exceeding this limit, random cropping is applied within each batch, and all sequences are padded to match the length of the longest sequence in the batch.

#### Masked Nucleotide (MN) Prediction

In the pretraining phase, we adopt a masked language modeling (MLM) strategy. Specifically, 25% of the nucleotide tokens are randomly masked, and the model is trained to predict the masked tokens based on the surrounding context. This approach facilitates the learning of contextual nucleotide representations.

Let *x* = (*x*_1_, *x*_2_, …, *x*_*T*_) denote a nucleotide sequence of length *T*, and let *M⊂* {1, 2, …, *T*} be the set of randomly selected masked indices, with |*M*| = 0.25*T*. The training objective for this task, referred to as masked nucleotide (MN) prediction, is:

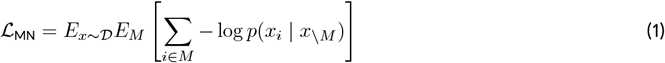

Here, *x*_*\M*_ denotes the unmasked tokens, and *p*(*x*_*i*_ | *x*_*\M*_) is the predicted probability of the true nucleotide at position

#### Secondary Structure (SS) Prediction

To integrate RNA structural features, we introduce a supervised learning task that leverages secondary structure (SS) annotations for the coding region, computed by ViennaRNA[36]. The SS is represented in dot-bracket notation: paired nucleotides are denoted by ‘(‘ and ‘)’, while unpaired nucleotides are denoted by ‘.’.

Using the same masking strategy as above, each nucleotide masked *x*_*i*_ is associated with a corresponding SS label *s*_*i*_, and the model is trained to predict this label. The SS prediction loss is defined as:

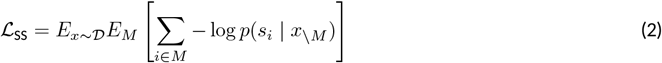

where *s*_*i*_ *∈* {(, ), .} represents the SS label at position *i*.

#### Total Objective

The overall training loss combines the MN and SS objectives with a weighting coefficient *λ*:

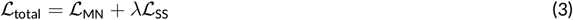

where *λ* balances the contribution of the SS task, with a default value of 0.4.

#### Training Details

Pretraining was performed on four NVIDIA V100 GPUs for 5 epochs, requiring approximately 10 days. The AdamW optimizer was used, with a batch size of 3 and a gradient accumulation step of 6. We reserved 0.3% of the training data for validation. The learning rate was linearly warmed up from 0 to 2e-4 over the first 10% of steps and then decayed to 0 following a cosine schedule.

Fine-tuning was performed using training data from 8 representative species, maintaining the same hyperparameters as pretraining. This phase utilized two NVIDIA V100 GPUs and ran for 10 epochs.

#### Model inference

During inference, the nucleotides corresponding to the known amino acid sequence are explicitly provided, while unknown regions are marked with the <unk> token. For example, for the amino acid sequence “MKD”, the input DNA sequence might be ATGAA<unk>GA<unk>. This approach, compared to using only the amino acid sequence, supplies richer contextual information, allowing the model to more effectively generate optimized DNA sequences. Alternatively, a complete and valid DNA sequence can be directly input for further optimization.

### 5.4 Model evaluation

To comprehensively evaluate the quality of the generated sequences, we utilized a variety of assessment metrics, primarily adapted from those employed in CodonTransformer[21]. Each metric focuses on a distinct aspect of codon usage or sequence characteristics. These evaluations are designed to assess the extent to which the generated DNA sequences align with natural sequences, considering both their biological relevance and structural properties.

Specifically, we evaluated:

- **Codon Recovery Rate**: Measures the percentage of exact codon matches between the generated and natural DNA sequences. A higher recovery rate indicates better preservation of native sequence structure and codon usage. For a natural DNA sequence *A* and a generated sequence *B*, the recovery rate is calculated as:

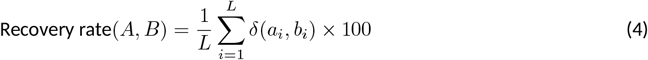

where *L* is the length of the sequences, *a*_*i*_ and *b*_*i*_ are the codons at position *i* in sequences *A* and *B*, respectively, and *δ* is the Kronecker delta function, which equals 1 if *a*_*i*_ = *b*_*i*_ and 0 otherwise.
- **Codon Similarity Index (CSI)**: The CSI[33], derived from the CAI[9], quantifies the similarity in codon usage between a sequence and the codon usage table of an organism, without relying on any highly expressed genes. CSI provides a softer and potentially more biologically appropriate estimator of codon usage for a given organism. The relative adaptiveness of a codon *w*_*ij*_ is calculated as the ratio of its frequency *x*_*ij*_ to that of the most frequently used synonymous codon 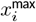 for the same amino acid:

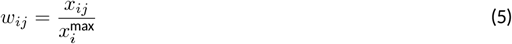

The CSI for a gene is then computed as the geometric mean of these relative adaptiveness values over all codons in a gene sequence of length *L*. A CSI value of 1 indicates that the sequence exclusively uses the most frequently used codons. Here, *w*_*k*_ denotes the value of *w*_*ij*_ corresponding to the current position *k*.

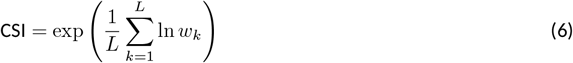
- **Codon Frequency Distribution (CFD)**: The Codon Frequency Distribution (CFD)[35] indicates the frequency of rare codons selected by the model (with a frequency *<* 0.3), which tends to vary across species. A weight *w*_*ij*_ is assigned to each codon as:

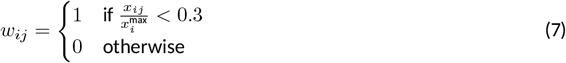

The CFD is then computed as the mean of these weights across all *L* codons in the sequence:

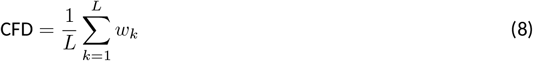
- **Low Frequency Codon (LFC)**: A codon is defined as a low frequency codon (LFC) for a given amino acid if its normalized selection frequency falls below 0.3 within the host organism’s genomic context. The binary identification function *I*_*ij*_ for codon *j* encoding amino acid *i* is formalized as:

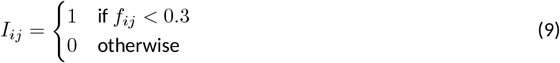

where *f*_*ij*_ represents the frequency of codon *j* for amino acid *i*.
- **GC Content (%GC)**: GC content[34] is a fundamental metric for measuring the proportion of guanine (G) and cytosine (C) in a DNA sequence. Different species and genes typically exhibit specific GC content ranges, which are closely related to gene stability and expression levels. It measures the proportion of guanine (G) and cytosine (C) bases in a DNA sequence.

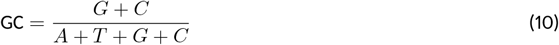
- **%MinMax**: The %MinMax[43] evaluates the balance between high-frequency and low-frequency codons within a sliding window along the sequence. This metric captures local fluctuations in codon usage, offering insights into the fine-grained distribution patterns that can influence translational efficiency and mRNA stability. For a window size *w* (typically *w* = 18), the %MinMax value at window *i* is computed as:

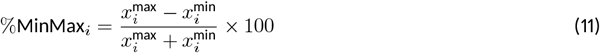

where 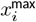 and 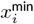 are the maximum and minimum codon usage frequencies within the *i*-th window, respectively.The overall %MinMax for the gene is an array of such values:

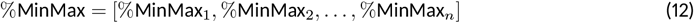

where *n* is the total number of windows in the sequence.
- **Dynamic Time Warping (DTW)**: Dynamic Time Warping (DTW)[37] is an algorithm designed to measure the similarity or alignment between two time series, particularly excelling in scenarios involving local distortions or shifts. In this study, we employ DTW to quantify the distance between the MinMax profiles of generated and native sequences, providing a more detailed evaluation of codon usage patterns across the entire sequence.The DTW distance between two time series *X* and *Y* is defined as:

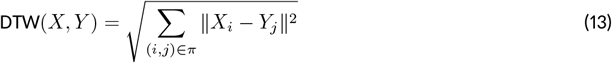

where *X* and *Y* are the %MinMax vectors, *π* is the optimal alignment path, and *X*_*i*_ *Y*_*j*_ denotes the Euclidean distance between aligned elements. DTW provides a robust quantitative assessment of how closely a generated sequence’s codon usage pattern follows the native profile. A smaller DTW distance indicates that the local dynamics of codon usage are more faithfully captured.
- **Minimum Free Energy (MFE) of RNA Structures**: Minimum Free Energy (MFE) reflects the stability of secondary structures (such as hairpin structures) formed by nucleic acid sequences at thermodynamic equilibrium. Lower energy generally indicates a more stable structure. Both overly stable and overly relaxed secondary structures can affect the transcription and translation efficiency of mRNA. RNA secondary structure prediction was performed using the fold() function from the ViennaRNA package[36], which computes the minimum free energy (MFE) of the resulting structures.

The code for calculating the Codon Recovery Rate, CSI, CFD, GC Content, %MinMax, DTW, and MFE is included in the source code. By incorporating these diverse metrics, we establish a comprehensive and multidimensional evaluation framework. This ensures that the model is not solely optimized for a single criterion (e.g., high CAI) but is also capable of capturing the subtle variations and biologically relevant features inherent in natural DNA sequences.

### 5.5 Model fitness score

To assess how well a coding DNA sequence aligns with the model’s learned codon preferences, we define a fitness score at the codon level. Given a codon sequence *c* = (*c*_1_, *c*_2_, …, *c*_*N*_) of length *N* (where each *c*_*i*_ represents a triplet codon), the model assigns a conditional probability *p*(*c*_*i*_ | *c*_*\i*_) to each codon based on its surrounding context.

The fitness score *F* (*c*) is computed as the average log-likelihood of the codons under the model:

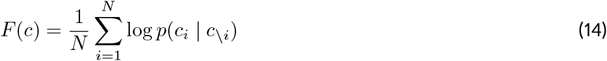

## 6 Acknowledgment

This work was partly supported by the National Key Research and Development Program of China under Grant No. 2024YFA0919702 (Y.W.) and 2023YFA0915500 (S.W. and L.Z.), National Science Foundation of China under grant no. 62272449 (Y.W.) and 12426303 (Y.W.), the Shenzhen Basic Research Fund under grant no KQTD20200820113106007, ZDSYS20220422103800001. We would also like to thank the funding support by the Key Laboratory of Quantitative Synthetic Biology, Chinese Academy of Sciences under grant no. CKL075. This research is also supported by the Science and Technology Development Fund of Macau (0004/2025/RIA1 to J.G.).

## 7 Data availability

The dataset used in this study consists of pretraining data, fine-tuning data, and a held-out test set. All processed sequences and related annotation files are publicly available at https://drive.google.com/drive/folders. The pretrained model weights can be downloaded from https://huggingface.co/guyuehuo/TransCodon.

## 8 Code availability

The source code used to train and evaluate the *TransCodon* model is publicly available at https://github.com/guyuehuo/TransCodon. The repository contains scripts for data preprocessing, model training, inference, and evaluation, along with detailed instructions for reproducing the results presented in this paper.

