## Supporting Materials for "Learning the native-like codons with a 5’UTR and RNA secondary structure aided species-informed transformer model"

### Part 1 Ablation Study: Effect of Vocabulary Granularity

A critical design choice in BERT-based modeling of DNA/RNA sequences is how to define the vocabulary that bridges nucleotide sequences and amino acid semantics. We compared two strategies:

- **Codon-level vocabulary:** Each codon (a triplet of nucleotides) is treated as a distinct token, jointly representing the codon and its corresponding amino acid. This is consistent with the strategy used by *CodonTransformer*[1].
- **Nucleotide-level vocabulary:** Each nucleotide (A, C, G, T) is treated as an individual token, aligning with the representation schemes adopted by most RNA/DNA pretrained models.

To assess the impact of vocabulary granularity, we conducted a controlled ablation experiment using **Dataset1**, with a training set identical to that of *CodonTransformer*. For testing, we randomly selected 50 sequences from the top 10% based on CSI across five species (*Arabidopsis thaliana, Escherichia coli, Homo sapiens, Mus musculus, and Saccharomyces cerevisiae*). This resulted in a total of 250 sequences for evaluation.

We evaluated performance using the **codon recovery rate**, a direct and interpretable measure of how effectively the original codons can be reconstructed from the model's output. As shown in Supplementary Table S1, models utilizing codon-level vocabulary (e.g., *TransCodon*, *CodonTransformer*) produced similar recovery scores. In contrast, models using nucleotide-level vocabulary demonstrated an average improvement of **2%** in codon recovery rate.

Further analysis revealed that sequences generated using the triplet-based vocabulary exhibited greater deviations from natural sequences in terms of minimum free energy (**MFE**) and **GC content**. Specifically, the average deviations for the single-nucleotide vocabulary were approximately 50 (MFE) and 3.2% (GC content), while for the triplet-based vocabulary, these deviations increased to approximately 90 and 4.2%, respectively.

Table S1: Average codon recovery rates and biological Deviation

| **Model** | **Codon Recovery Rate (%)** | **Biological Deviation** | |
|---|---|---|---|
| | | **Avg. MFE Deviation ↓** | **Avg. GC Content Deviation ↓** |
| TransCodon (Codon-level) | 54.5 | ~90 | ~4.2% |
| TransCodon (Nucleotide-level) | **56.5** | **~50** | **~3.2%** |
| CodonTransformer | 54.1 | ~65 | ~5.2% |

These results suggest that finer-grained nucleotide representations yield sequences that are both more accurately recoverable and more biologically plausible. Consequently, we adopt the nucleotide-level vocabulary as the default for our model.

### Part 2 Ablation Study:Exploration of Training Data Volume

In the era of large-scale models and data, it is widely recognized that increasing both the training data volume and model parameters often leads to improved performance. Therefore, we explored the impact of model capacity and training data volume on the performance of our framework. Existing models, such as CaLM[2] and CodonTransformer[1], are typically built with around 90 million parameters, whereas PlantRNA-FM[3] adopts a smaller configuration with 35 million parameters, balancing training costs and generalization ability. In line with this, we selected a similar parameter scale of approximately 90 million for TransCodon as a reasonable reference point.

Unlike general-purpose pretraining models such as CaLM or PlantRNA-FM, which aim to learn biologically meaningful representations of DNA/RNA language, the primary objective of TransCodon is to optimize codon usage across species. For language models, expanding the training data typically leads to direct improvements in performance. However, TransCodon faces a different challenge: it is constrained by the limited number of DNA sequences available for each species. As more data is added, it primarily introduces new codon distributions from additional species, raising the question of whether this expansion enhances the codon landscape of previously encountered species.

To assess the impact of training data scale, we conducted an ablation study by progressively increasing the size of the training dataset and evaluating model performance on a fixed test set. All test species were selected from **Dataset1** (consistent with the main text), ensuring no overlap with species from Datasets 2, 3, or 4. We used **codon recovery rate** and the average Dynamic Time Warping (**DTW**) distance between the generated and natural sequences as evaluation metrics.

As shown in Figure S1a, increasing the training data from 1M to approximately 5.5M resulted in little change in both the codon recovery rate and DTW distance. This suggests that the model performs stably within this range, effectively incorporating codon distribution patterns from a broader range of species. However, when the training data was expanded to 11M, both evaluation metrics worsened, indicating that this data scale may surpass the modeling capacity of the current model size.

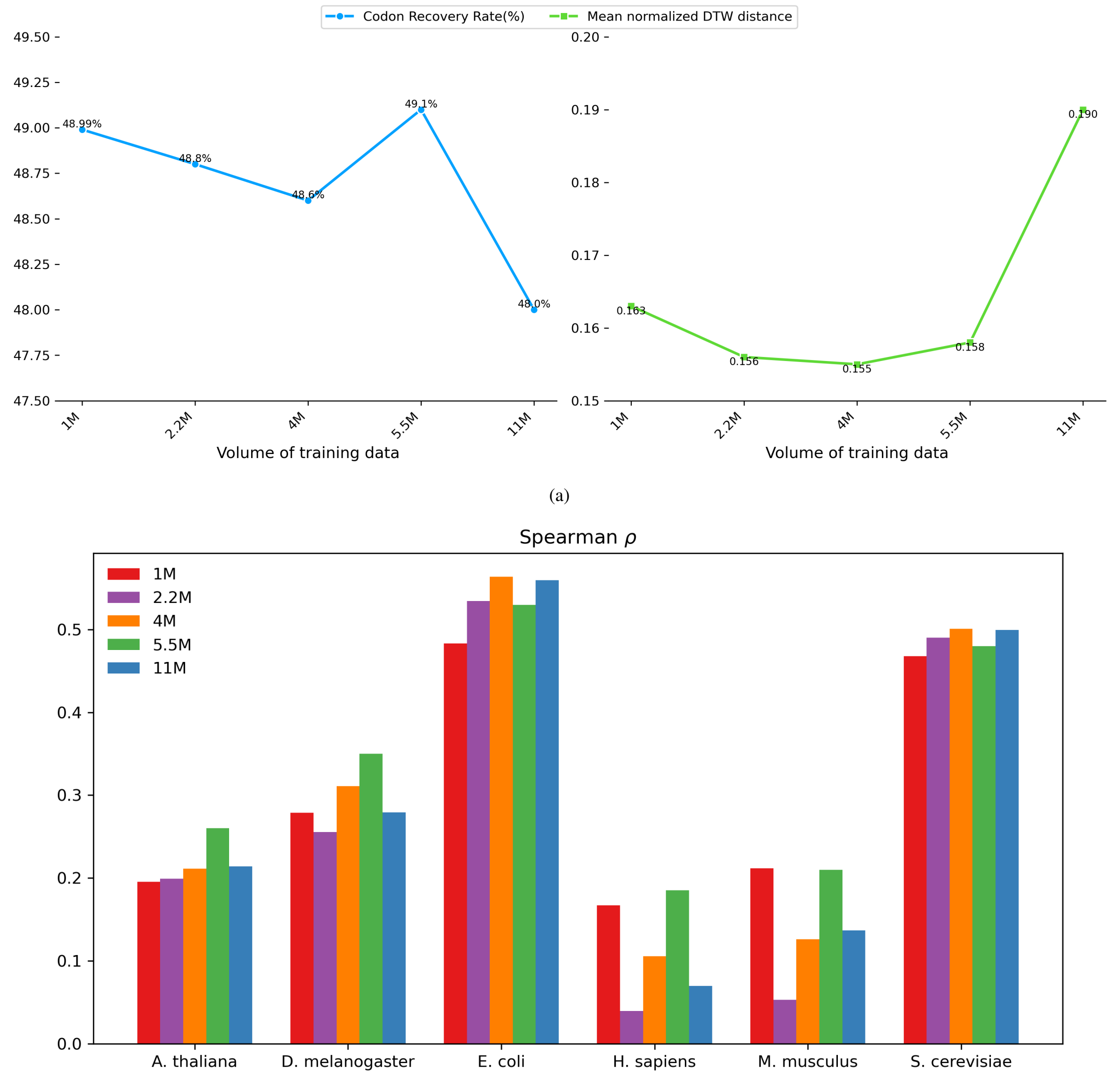

(a)

(b)

Figure S1: Exploration of Training Data Volume. Specifically, the training sets labeled 1M, 2.2M, 4M, and 5.5M correspond to Dataset1, Dataset1+Dataset2, Dataset1+Dataset2+Dataset3, and Dataset1+Dataset2+Dataset4, respectively. The 11M training set consists of Dataset1+Dataset2 combined with 8.8M bacterial data points. **(a)** Average codon recovery rate and average DTW distance across 250 sequences for different training data configurations. **(b)** Correlation between model-predicted fitness score and protein abundance across six species for different training datasets.

Furthermore, although the test set is composed solely of species from **Dataset1**, we conducted a detailed analysis of the six species discussed in the main text to investigate how the correlation between predicted fitness and protein abundance (from PaxDb[4]) evolves with increasing training data size. As shown in Figure S1, there is a general upward trend in the Spearman correlation as the training data increases, although the improvement remains relatively modest.

Interestingly, we observed that the correlation varies across species and appears to be closely tied to the composition of the training data. For complex eukaryotic organisms such as *Arabidopsis thaliana*, *Drosophila melanogaster*, *Homo sapiens*, and *Mus musculus*, the correlation decreased when the training data increased from 1M to 2.2M, with the exception of *A. thaliana*. This decline is likely due to the introduction of **Dataset2**, which consists solely of archaeal species, introducing biases that misalign with the regulatory features of higher eukaryotes. However, as the training data expanded to 5.5M, incorporating fungal sequences from **Dataset4**, the correlations for all four species reached their peak. In contrast, when the dataset grew to 11M without the addition of further fungal data, performance deteriorated once again.

For microbial species such as *Escherichia coli* and *Saccharomyces cerevisiae*, the inclusion of **Dataset2** (archaea) and **Dataset3** (bacteria) led to an improvement in correlation. However, incorporating **Dataset4** (fungi) in the 5.5M setting caused a slight decline. These findings suggest that even when the test set remains fixed, model performance is highly sensitive to both the quantity and the taxonomic composition of the training data.

In summary, increasing the training data from 1M to 5.5M primarily enabled the model to more effectively capture codon usage patterns across diverse species. These results underscore the importance of training data diversity for model generalization and suggest a key direction for future work: expanding model capacity to better learn from larger, more heterogeneous training datasets.

### Part 3 Ablation Study:Evaluating the Effect of Adding 5' UTR to Training Data

In the previous experiments, the test set was drawn from Dataset1, where the training data did not include 5' UTR regions. To evaluate the impact of incorporating 5' UTR information, we trained a new model using a combined dataset (Dataset1 + Dataset2), where Dataset2 is an archaeal dataset containing 100 bp of annotated 5' UTR sequences. For comparison, we also trained a baseline model without any 5' UTR input. The test set comprised 8 species from **Dataset2** (*Halobacterium salinarum, Halobacterium litoreum, Thermococcus celer, Pyrobaculum calidifontis, Methanococcus vannielii, Methanosarcina mazei, Desulfurococcus amylolyticus, and Picrophilus oshimae*), ensuring no overlap between training and testing samples.

We compared the performance of two models—one trained with 5' UTR input and one without—based on the average codon recovery rate across the eight species. As shown in Figure S2, even without 5' UTR data, the baseline model achieves competitive performance. When the model trained with 5' UTR data is tested using only amino acid sequences (i.e., without 5' UTR input during inference), its performance is slightly inferior to the baseline. However, when the full 5' UTR sequence is provided during inference, the model demonstrates improved codon recovery performance.

Additionally, we assessed the model's fitness correlation with protein abundance for *Halobacterium salinarum* using data from PaxDb. The results were consistent with previous findings: while incorporating 5' UTR sequences enhances model accuracy, the model requires 5' UTR input during inference for optimal performance. Without this input, performance may degrade compared to models trained exclusively on coding sequences.

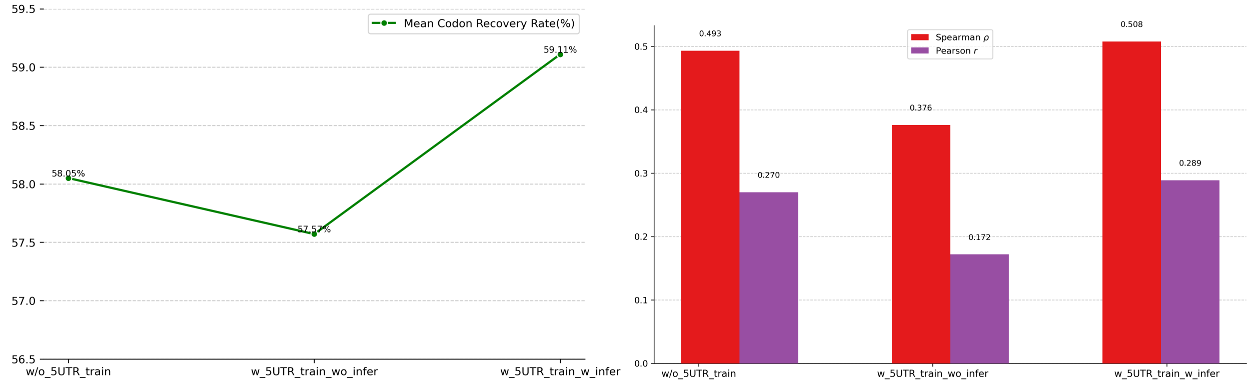

Figure S2: Evaluating the Effect of Adding 5' UTR to Training Data. In detail,the notations `w/o_5UTR_train`, `w_5UTR_train_wo_infer`, and `w_5UTR_train_w_infer` denote training without 5' UTR involvement, training with 5' UTR but inference without 5' UTR, and both training and inference with 5' UTR participation, respectively.

### Part 4 Ablation Study:Effect of RNA Secondary Structure Supervision on Training

To explore whether incorporating RNA secondary structure as an auxiliary supervision signal could enhance model performance, we conducted an ablation study using the same training setup as the 5' UTR experiment. Specifically, we utilized a combined dataset consisting of Dataset1 and Dataset2. Two models with identical architectures were trained: one using only masked language modeling (self-supervised learning) and the other incorporating an additional supervised objective to predict RNA secondary structure. Evaluation was performed on eight held-out species from Dataset1, consistent with the main test set, ensuring no overlap between training and testing data. Performance was assessed using two metrics: the average Dynamic Time Warping (DTW) distance between generated and natural sequences, and the correlation between model-inferred fitness scores and protein abundance from PaxDb across six species.

For the average DTW distance across the eight test species, the model incorporating RNA secondary structure supervision achieved a score of 0.156, compared to 0.174 for the model trained solely with the masked language modeling objective. Furthermore, Figure S3 shows that incorporating RNA secondary structure supervision improves the correlation between predicted fitness scores and protein abundance across most species, with the exception of *Escherichia coli*. These results suggest that incorporating RNA structural information as a supervisory signal provides modest but consistent improvements across both evaluation metrics. While the performance gain is not substantial, the findings indicate that RNA secondary structure acts as a useful inductive bias, helping the model learn sequence representations that better capture the underlying biological patterns.

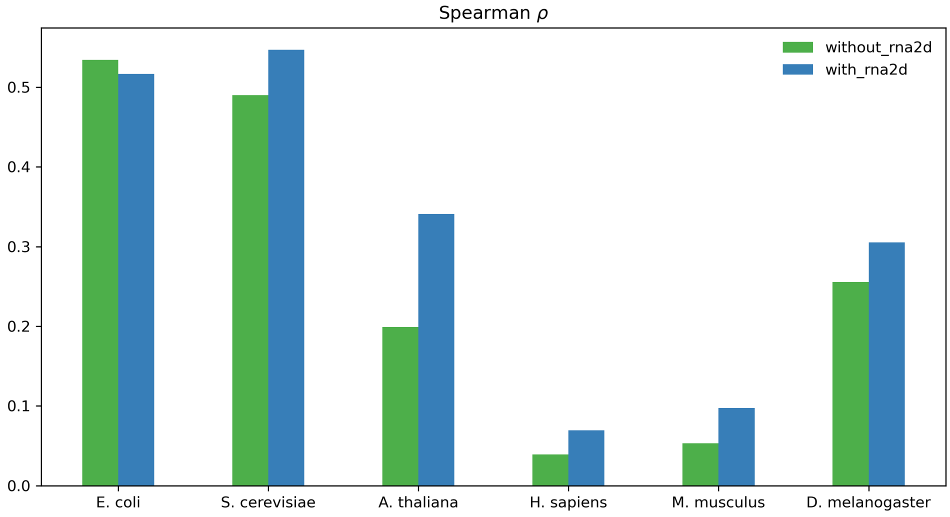

Figure S3: Evaluating the Impact of RNA Secondary Structure Supervision on Training. The notations `with`$_rna2dand$

### Part 5 Exploration of Incorporating Embeddings from RNA/DNA Large Language Models

With the rapid emergence of large-scale RNA/DNA pre-trained models, such as evo2[5] and RNA-FM[6], both model size and training data have expanded dramatically—RNA-FM, for example, was trained on over 23.7 million sequences, far surpassing the scale of our dataset. This raises an intriguing question: can the embeddings from these larger models be integrated as supplementary information to enhance our model's ability to capture the deeper biological insights encoded in DNA? This represents a promising avenue for future exploration.

Specifically, we first experimented by feeding natural DNA sequences into RNA-FM to extract embeddings, which were then used as additional input features during model training. This approach led to a modest improvement in performance on the codon recovery task: the average recovery rate across eight test species from Dataset1 increased from 49% to approximately 51%. This improvement is expected, as the embeddings were derived from natural DNA sequences. However, a more challenging and meaningful scenario lies in determining whether such external embeddings can enhance performance without direct access to the native DNA sequence—i.e., through an iterative inference strategy.

In this iterative setup, we begin with a random input amino acid sequence to generate a predicted DNA sequence (seq1). We then use seq1 to obtain an RNA-FM embedding, which is subsequently input to generate a refined sequence (seq2). This process is repeated in successive iterations to refine the output. Despite conducting three rounds of such iterations, no significant improvement in codon recovery was observed. This lack of improvement is likely due to the noise introduced by using imperfect input sequences in RNA-FM, which leads to biased embeddings. Therefore, while incorporating external embeddings from large RNA/DNA language models is conceptually appealing, further work is necessary to effectively harness these embeddings in practice.

### Part 6 Supplementary Figure

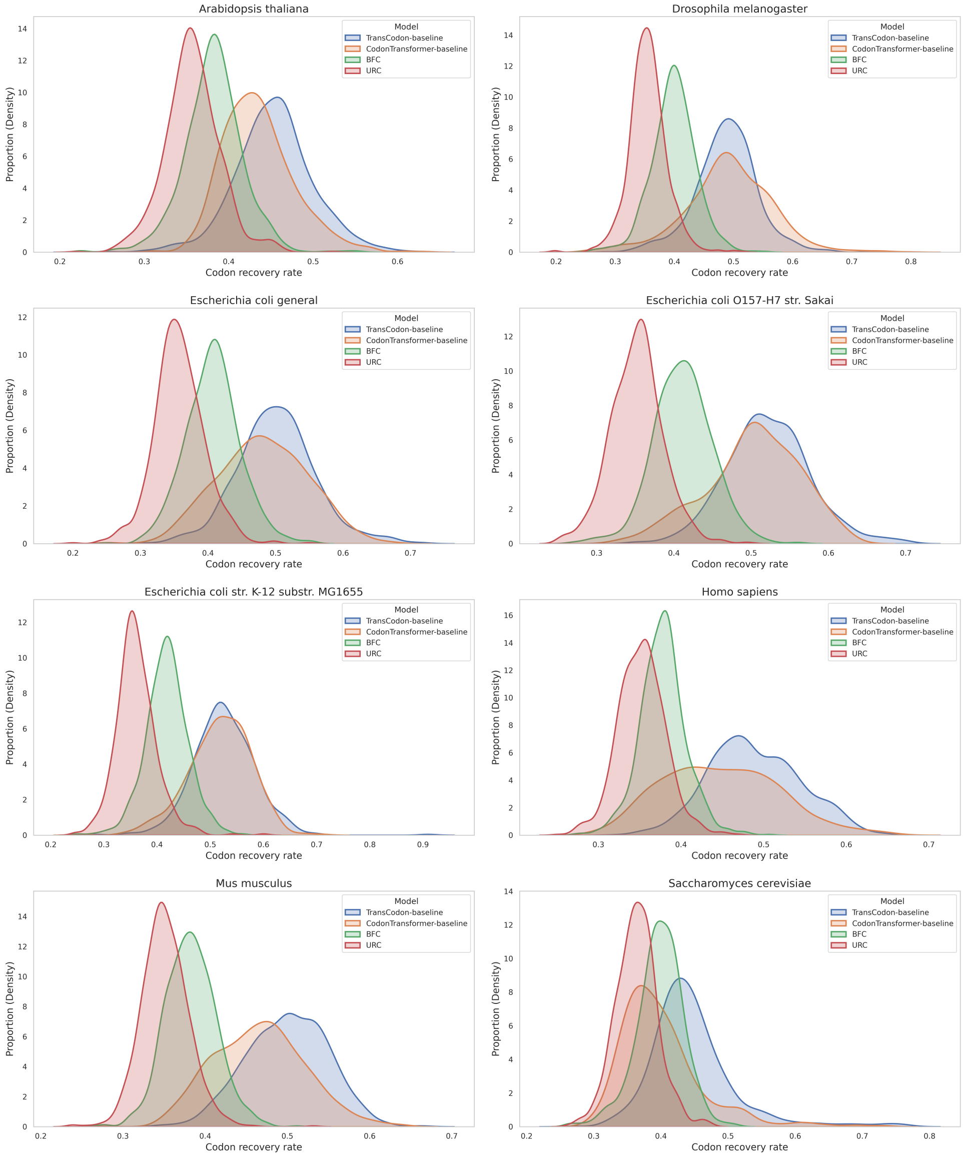

Figure S4: TransCodon-generated sequences exhibit a close resemblance to natural ones. The codon recovery rates of various methods across all 8 species.

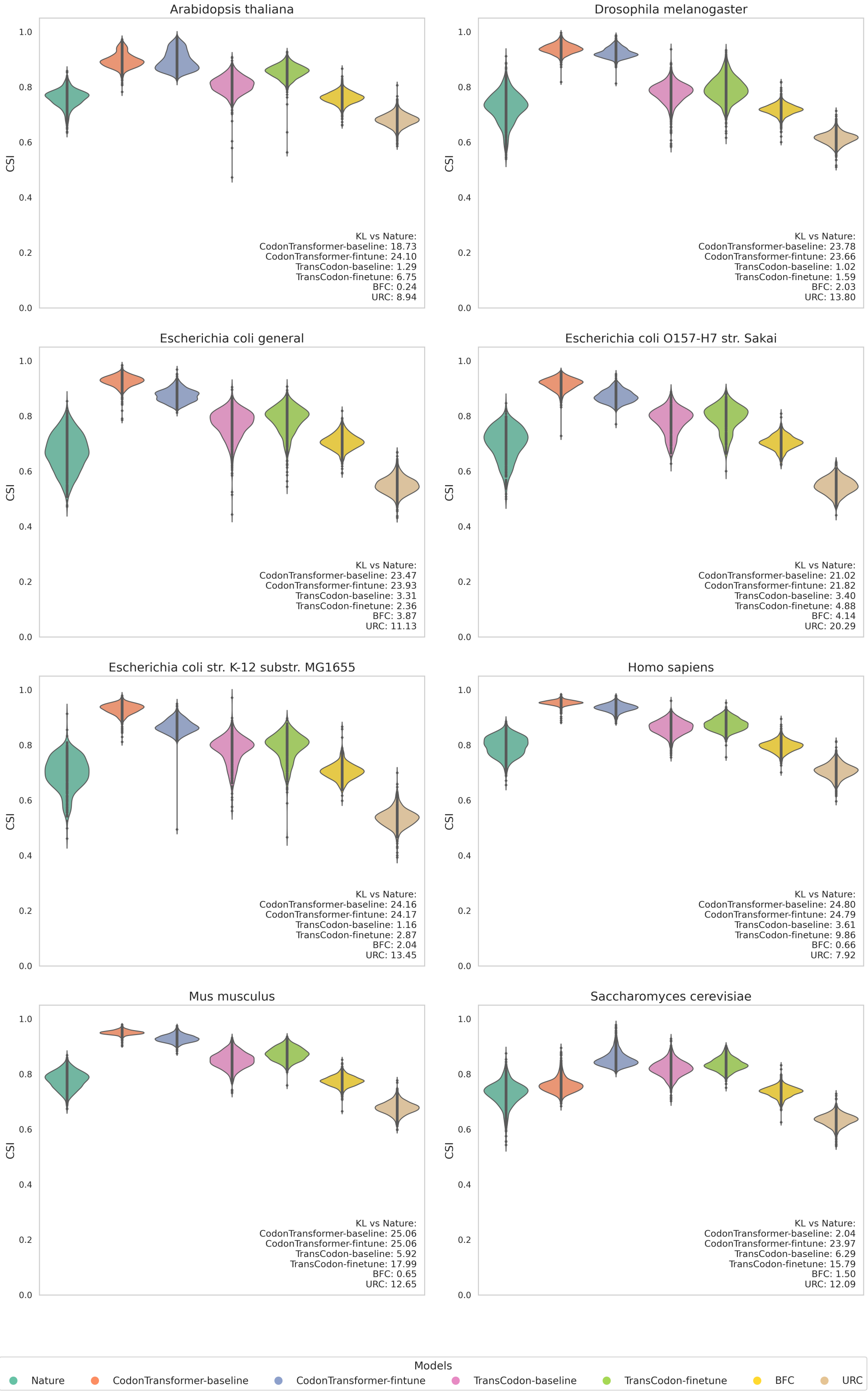

Figure S5: TransCodon-generated sequences exhibit a close resemblance to natural ones. The CAI of various methods across all 8 species.

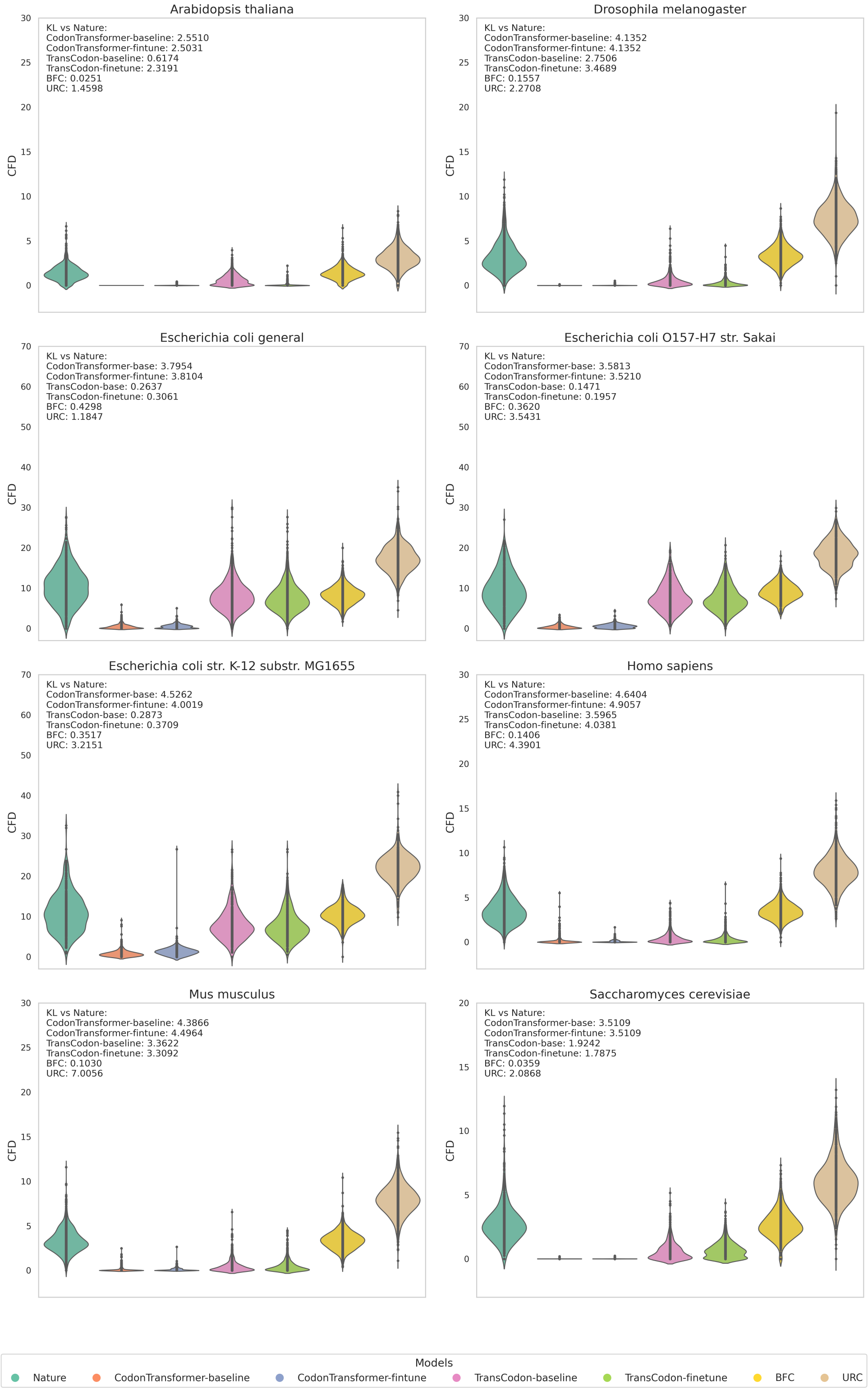

Figure S6: TransCodon-generated sequences exhibit a close resemblance to natural ones. The CFD of various methods across all 8 species.

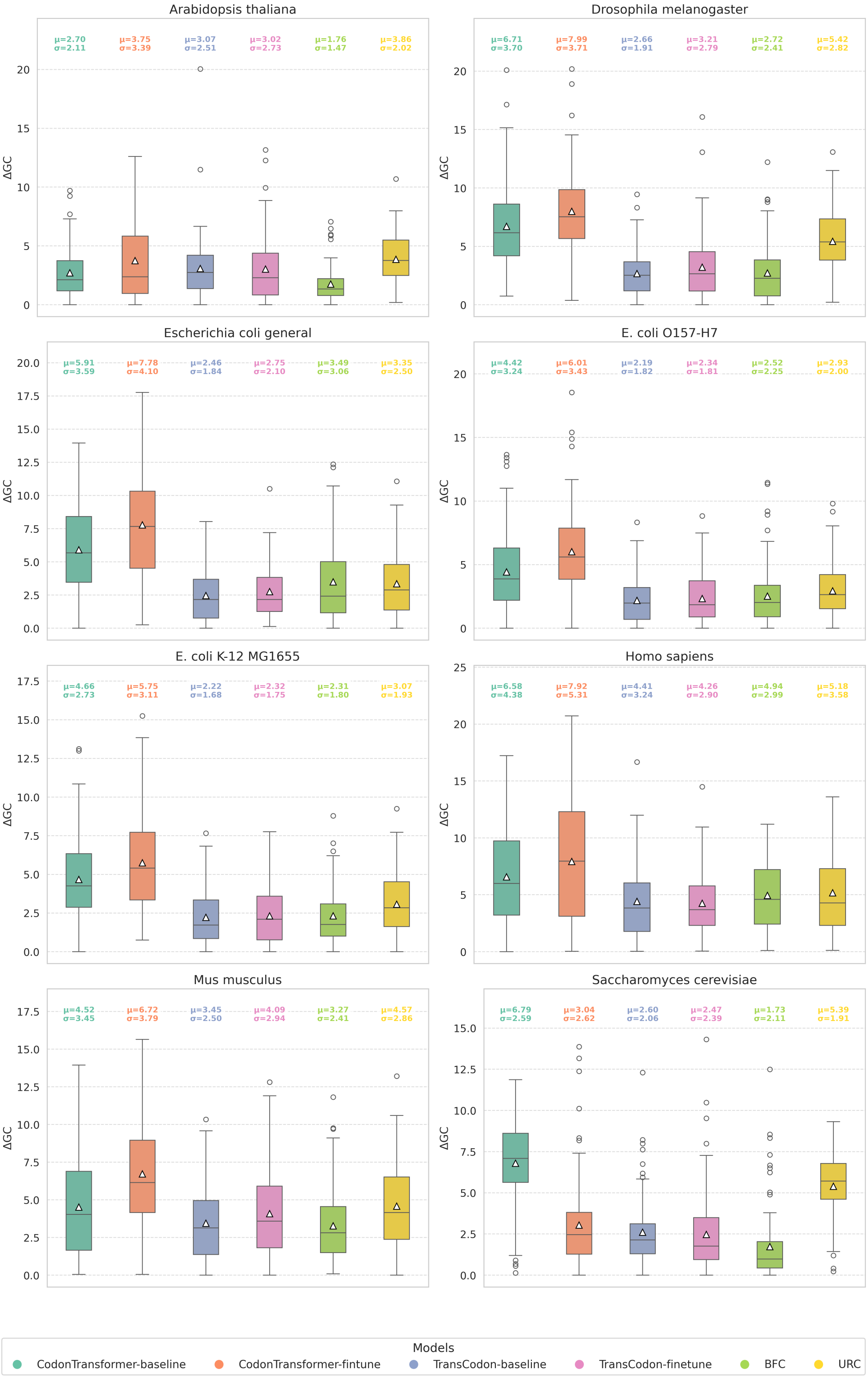

Figure S7: TransCodon-generated sequences exhibit a close resemblance to natural ones. The GC content deviation values of various methods across all 8 species.

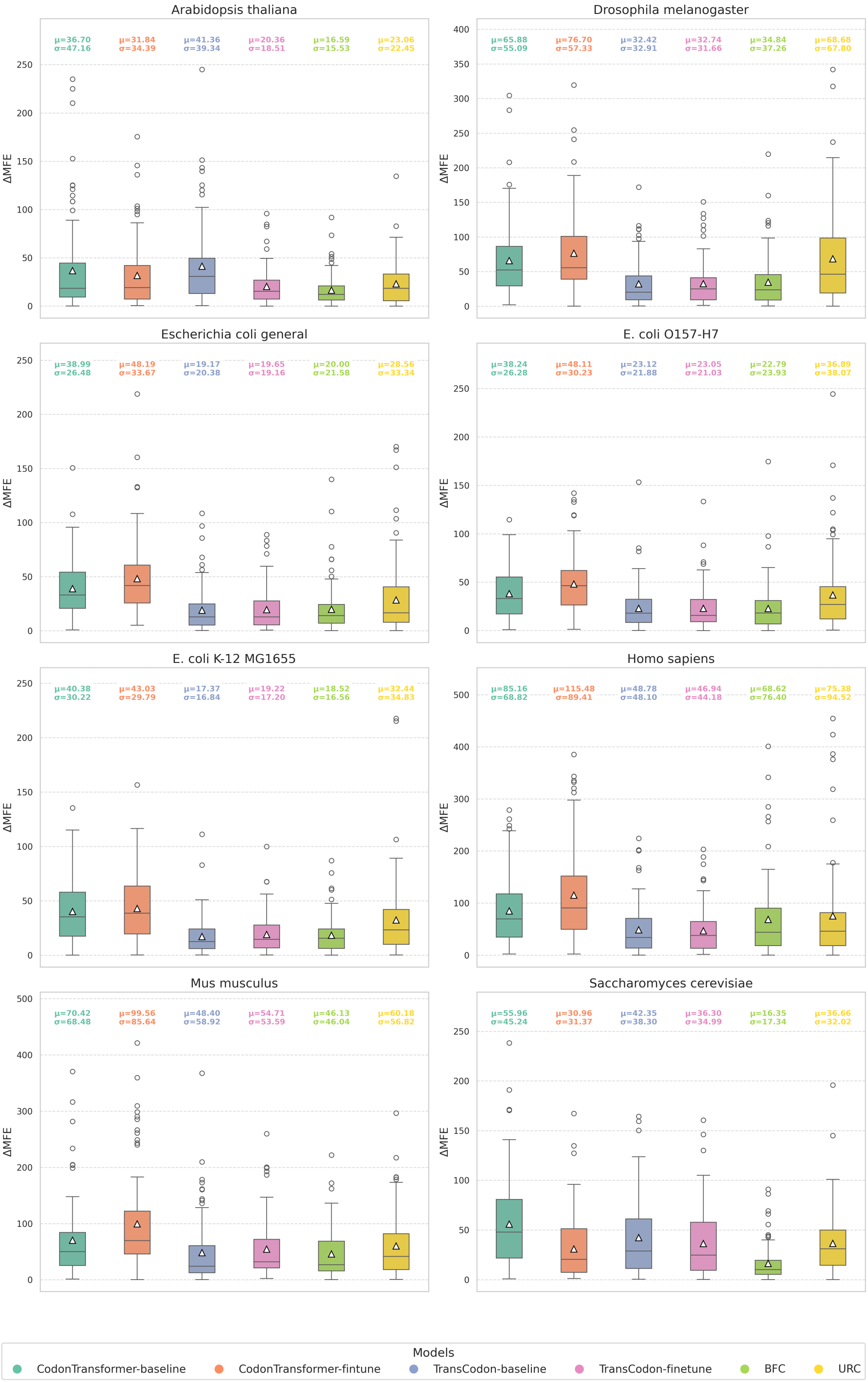

Figure S8: TransCodon-generated sequences exhibit a close resemblance to natural ones. The MFE deviation values of various methods across all 8 species.

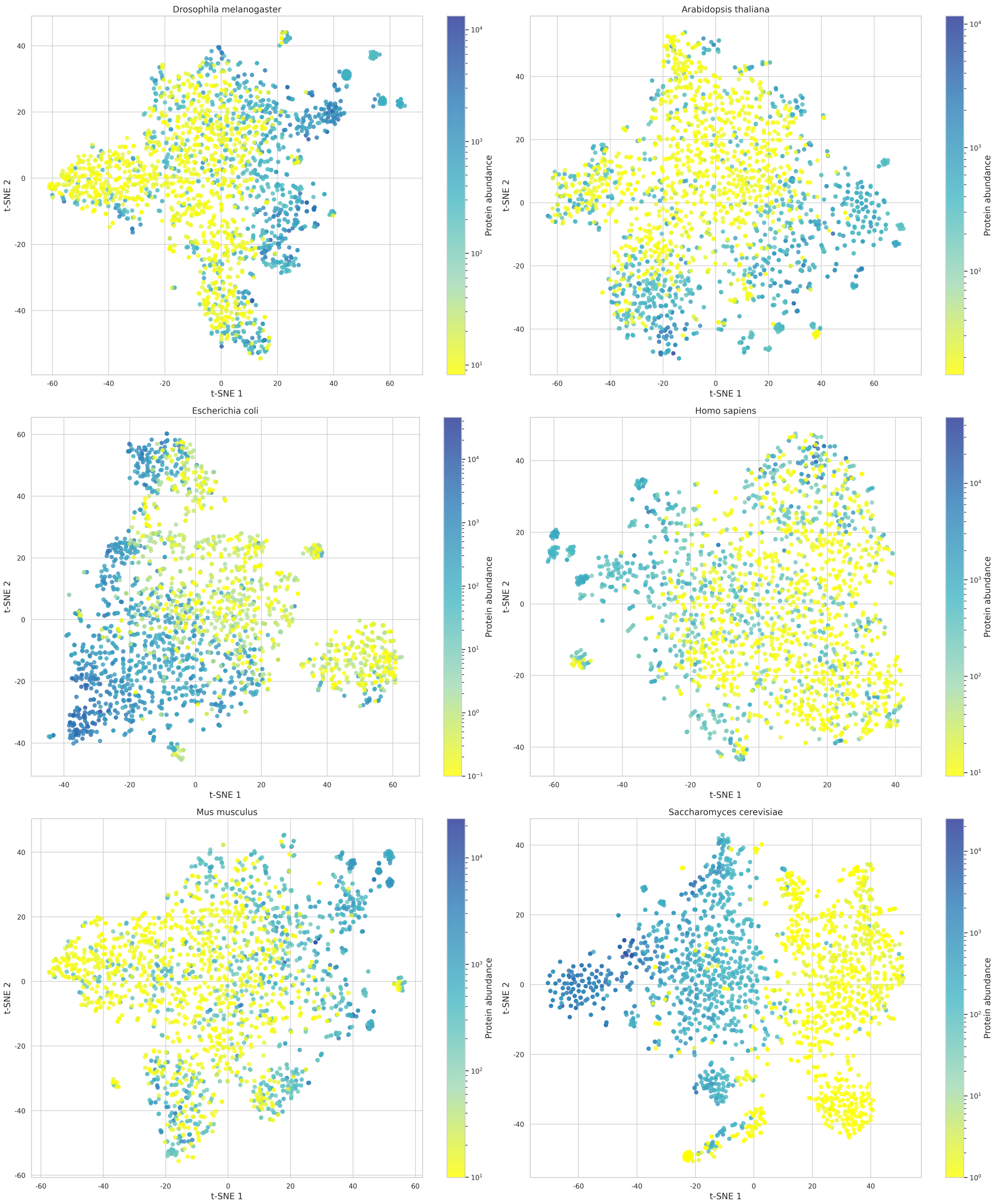

Figure S9: Dimensionality reduction of embeddings for the top 1,000 and bottom 1,000 protein abundance sequences for 6 species using TransCodon.

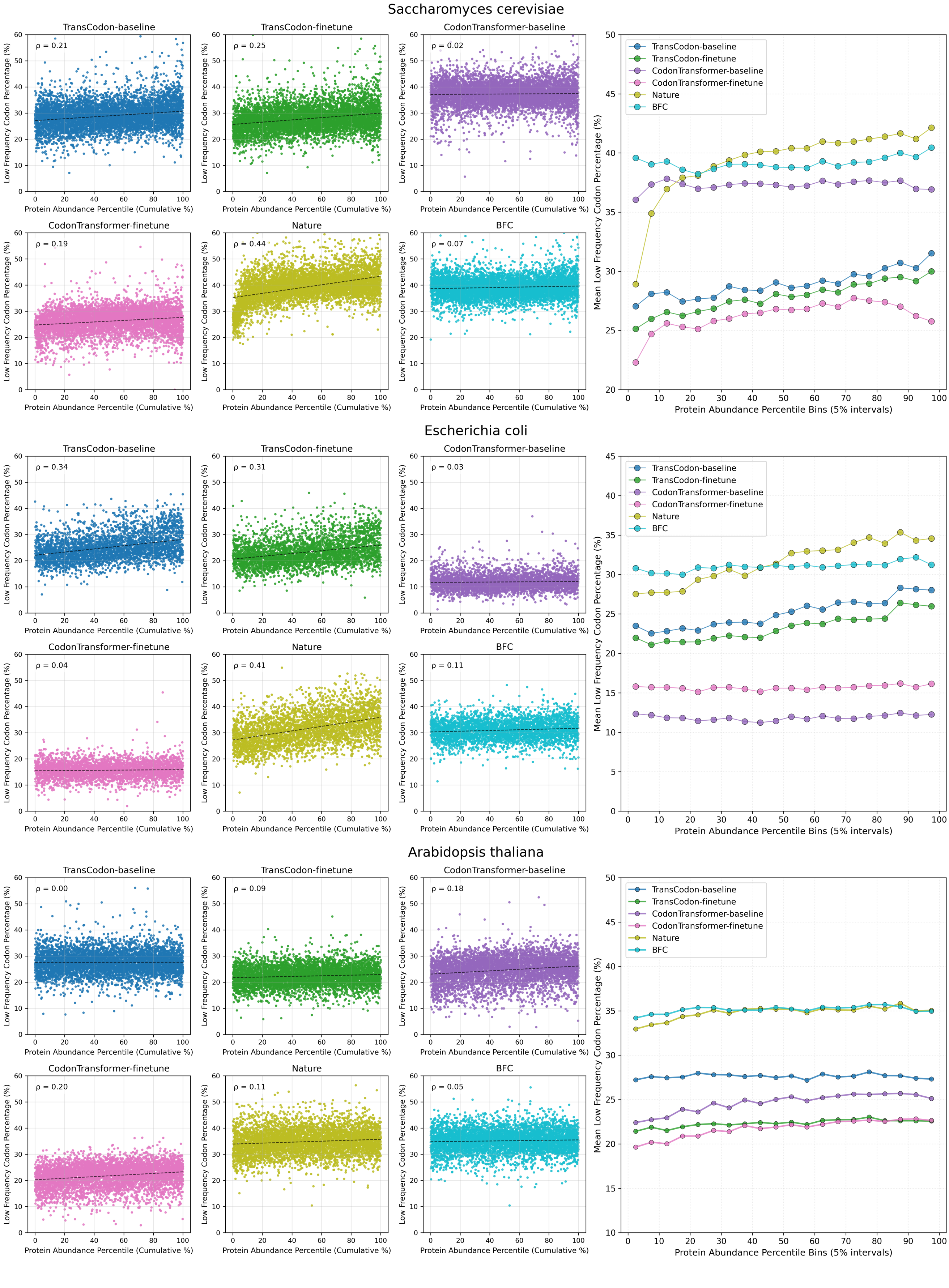

Figure S10: Illustrate the relationship between protein abundance and low-frequency codon usage for 3 species.

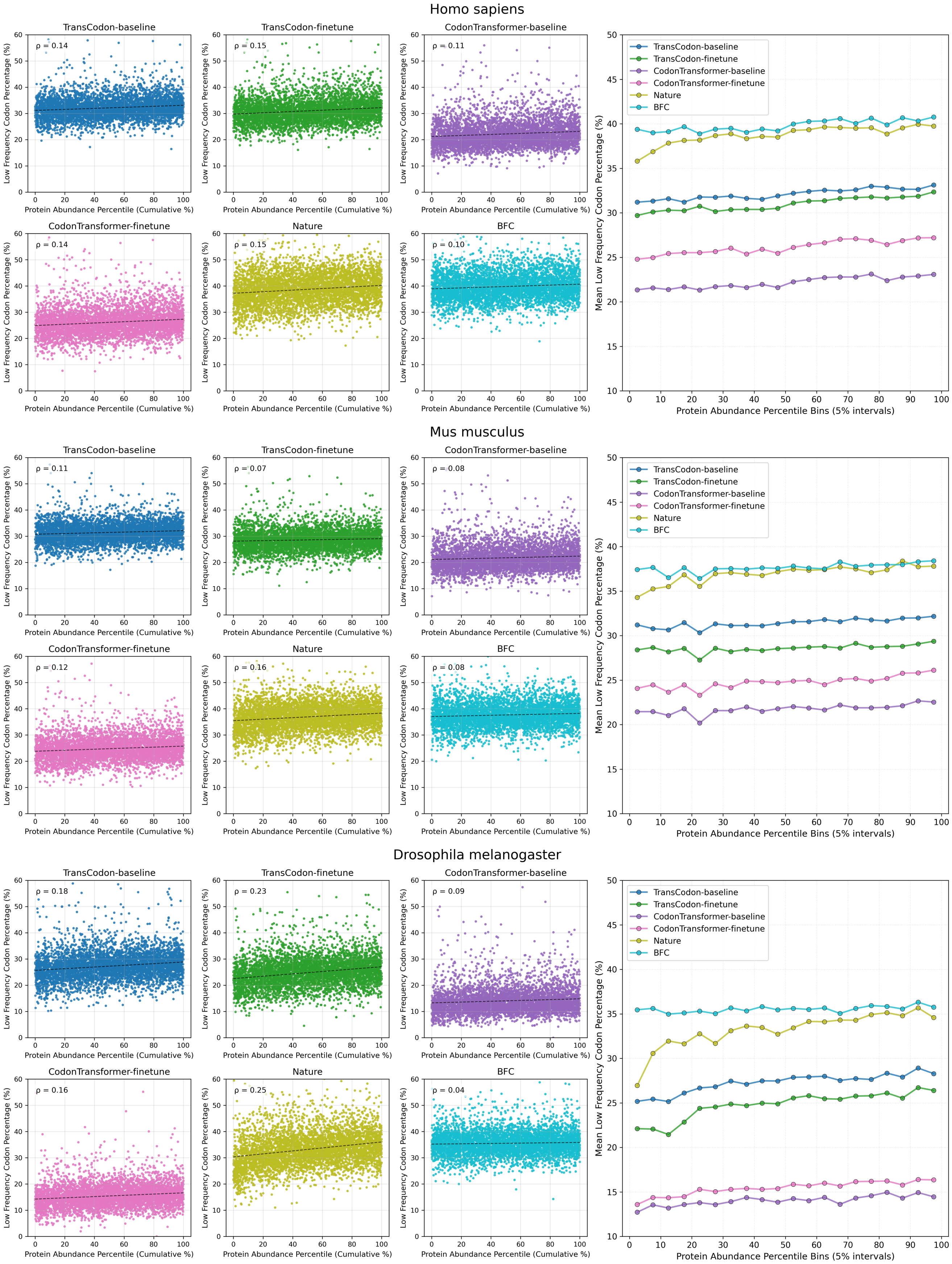

Figure S11: Illustrate the relationship between protein abundance and low-frequency codon usage for 3 species.
